# IsoMatch enables population-scale transcript integration and reveals reference-dependent isoform novelty

**DOI:** 10.64898/2026.10.09.758114

**Authors:** Xinchang Zheng, Sonia Garcia-Ruiz, Yilei Fu, Emil K. Gustavsson, Mina Ryten, Fritz J. Sedlazeck

## Abstract

Transcriptome analyses increasingly generate large collections of transcript models that must be integrated, compared and interpreted across samples and reference annotations. Transcript novelty depends on structural equivalence criteria and reference annotation. Here we present IsoMatch, a scalable framework that harmonizes transcript merging and reference-based classification through explicit, configurable structural criteria. IsoMatch preserves source-GTF and original-transcript traceability throughout the workflow, allowing every merged and classified model to be traced back to its contributing inputs. Benchmarking showed that IsoMatch preserves distinct transcript models during integration through configurable matching parameters for both splice junction chains and transcript ends. With its index-based architecture, IsoMatch merged 163 ENCODE transcript catalogs and classified 7.9 million transcript models with peak memory usage of 0.49 GB and 1.1 GB, respectively. Integrating GENCODE, RefSeq and CHESS revealed that 71.2% of transcript groups remained annotation-specific even without terminal constraints. For 4.84 million merged transcript models from 119 long-read transcriptomes from the 1000 Genomes Project, the integrated reference increased the full-splice match proportion from 12.0-13.6% with individual annotations to 18.0%. IsoMatch enables scalable transcript integration and classification with explicit structural criteria and preserved traceability, demonstrating how reference choice shapes estimates of transcript novelty.

## Introduction

Long-read RNA sequencing has transformed transcriptome analysis by enabling direct observation of full-length transcript structures(Ament et al. 2025; Monzó et al. 2025). Within single contiguous reads, it resolves exon connectivity, alternative splice junctions, transcript start sites (TSS), and transcript end sites (TES), allowing complex isoforms to be characterized directly rather than inferred through assembly. As long-read datasets grow, a large fraction of the transcripts are reported as novel(Patowary et al. 2024; Glinos et al. 2022). Several studies reported that around 30-70% of transcripts are outside of the reference annotations(Zheng et al. 2026; Clavell-Revelles et al. 2025; Glinos et al. 2022). A spread of that magnitude is unlikely to reflect biology alone. It also reflects how transcript models are compared, integrated, and interpreted after they are called(Pardo-Palacios, Wang, et al. 2024). Although these challenges have become particularly apparent with long-read RNA sequencing, they are not specific to a sequencing technology: transcript models generated by short-read assembly, hybrid approaches or reference annotation pipelines ultimately face the same problems of structural integration and classification (Kovaka et al. 2019; Prjibelski et al. 2023). These analytical choices become increasingly consequential as transcriptomic studies expand from individual samples to population-scale cohorts and as transcript models are compared across datasets and analysis workflows.

A fundamental decision in transcript integration is how structural equivalence between transcripts is defined. Two transcripts may share an identical splice-junction chain but differ slightly in their TSSs or TESs, while small offsets in junctions can also arise from variation in sequencing, alignment, or transcript calling. The tolerance applied to those differences determines whether two models are treated as the same or different transcripts. Permissive criteria can collapse biologically distinct isoforms, whereas overly stringent criteria can fragment equivalent transcript models because of minor coordinate differences. Before transcript novelty can therefore be assessed, transcript equivalence itself must first be defined. This definition should be explicit and configurable, with independent control over splice-junction, TSS, and TES matching rather than being implicitly determined by splice-junction agreement alone or tool-specific behavior. In practice, however, most existing tools rely almost exclusively on splice-junction agreement and disregard TSS and TES, or tie transcript equivalence to assumptions about a particular library preparation protocol (e.g., CapTrap), which limits their applicability across datasets(Kuo et al. 2020; M. Pertea et al. 2015; G. Pertea and Pertea 2020).

A second decision is which reference annotation defines whether a transcript is known or novel. Widely used annotations such as GENCODE(Mudge et al. 2025), RefSeq(Goldfarb et al. 2025), and CHESS(M. Pertea et al. 2018) describe overlapping but non-identical transcript sets. Consequently the same query transcript may be annotated as known in one annotation while being classified as novel against another. Transcript novelty is therefore a ‘moving’ target relative to the annotation against which it is evaluated rather than an intrinsic property of the transcript itself. Despite the widespread use of these reference resources, the extent to which their differences influence transcript classification has not been systematically quantified, and current workflows such as SQANTI3 are typically designed to compare query transcripts against one reference annotation at a time(Pardo-Palacios, Arzalluz-Luque, et al. 2024).

These issues are further complicated by the separation of transcript merging and reference-based classification in existing workflows. Transcript models are commonly first integrated across samples using tools such as GffCompare(G. Pertea and Pertea 2020) or TAMA(Kuo et al. 2020) and subsequently classified against a reference annotation using tools such as SQANTI3(Pardo-Palacios, Arzalluz-Luque, et al. 2024). Population-scale transcript merging and classification are therefore not simply repeated: the structural criteria used to construct the merged catalog directly determine which transcript models are subsequently interpreted against the reference. Once transcripts are merged across samples, preserving their relationships to original transcript identifiers and source samples also becomes essential for interpreting the resulting catalog. Existing merging tools commonly provide such information through separate tracking files, which can complicate filtering, propagation, and downstream analysis when hundreds of transcript sets and millions of transcript models are involved. At this scale, computational efficiency also becomes a practical requirement. Together, these considerations expose three practical limitations of current large-scale transcript workflows: limited scalability across large GTF collections, loss or separation of transcript traceability during integration, and disconnected integration and classification steps that complicate downstream interpretation.

IsoMatch was developed to address these three limitations within a unified GTF-level framework. It scales transcript integration and classification to millions of transcript models, retains the identity and source of every contributing transcript, and uses a common structural representation of splice junctions, TSSs and TESs across merging and reference-based interpretation. Transcript equivalence is explicitly defined through independently configurable matching of splice-junction chains, TSSs, and TESs separately, including user-defined tolerance for each coordinate offsets. This allows experimental or technical wobble to be accommodated without implicitly treating it as transcript novelty, while permitting the required stringency to be matched to the experimental design. The resulting merged transcripts retain their source samples and original transcript identifiers, enabling traceability to be preserved throughout downstream analysis. The same framework provides reference-based classification with SQANTI3-compatible structural categories, allowing merged transcript catalogs to be directly interpreted against individual or unified reference annotations. An indexed implementation enables both merging and classification to scale to millions of transcript models with low memory requirements.

In this study, we systematically benchmarked IsoMatch against established transcript merging and classification tools and showed that it preserves transcript structure and achieves high classification concordance while substantially reducing computational requirements. IsoMatch scaled to hundreds of cohort-level GTF files and millions of transcript models with low memory usage, and was further stress-tested with 1,000 GTF inputs. We further used IsoMatch to systematically compare GENCODE (v49 basic)(Mudge et al. 2025), RefSeq (v110)(Goldfarb et al. 2025), and CHESS (v3.1.3)(M. Pertea et al. 2018) revealing substantial differences among these widely used reference annotations. Only 7.74% of merged transcript models were shared across all three annotations, and even when transcript boundaries were ignored, 71.18% remained specific to a single annotation. Changing the structural equivalence criteria further altered the extent of cross-annotation overlap, demonstrating that transcript integration depends directly on how equivalence is defined. Finally, integrating the three reference annotations increased the full-splice match (FSM) proportion of 4.84 million population-level transcripts from 12.0-13.6% with individual references to 18.0%, showing that reference choice substantially influences estimates of transcript novelty. Together, these results establish IsoMatch as a scalable framework for making transcript equivalence and reference context explicit in population-scale transcript integration and classification.

## Results

### IsoMatch connects transcript merging and reference-based classification in a unified workflow

IsoMatch consists of three modules: index, merge, and classify that form a unified framework for harmonized transcript integration and reference-based classification (**Figure 1A**). Both the merge and classify functions in IsoMatch operate on indexes built from individual GTF files. First, each single-sample GTF is indexed to generate a compact index file (.isomx). This file stores isoform structures and splice-site sequence motifs, the latter requiring a reference genome file. A separate file (.isoms) is also generated to store transcript-level attributes (e.g. transcript_id) (**Figure 1B**). Second, all indexed GTF files are merged into a unified transcript catalog. During merging, IsoMatch follows a junction-chain-prioritized strategy while also considering TSS and TES as merge criteria (**Methods**). Specifically, transcripts from all input GTF files are first grouped by strand and junction chain. Transcripts are classified by splice-site motif, with GT-AG, GC-AG and AT-AC treated as canonical, consistent with SQANTI3(Pardo-Palacios, Arzalluz-Luque, et al. 2024); a transcript is considered non-canonical if any of its junctions falls outside these three motifs. Junction matching can optionally allow wobble to absorb small splice-site coordinate differences, applied separately to the two groups (default 0 bp for canonical transcripts and 3 bp for non-canonical transcripts). The grouped transcripts are then further split according to user-defined TSS and TES distance thresholds (default 50 bp). Finally, the splice-junction chain, TSS, and TES of the representative transcript are selected from those observed among the source transcripts. During this step, IsoMatch optionally accepts independent TSS and TES evidence regions to inform the selection of the representative transcript boundaries (**Figure 1C** and **Methods**).

**Figure 1.**
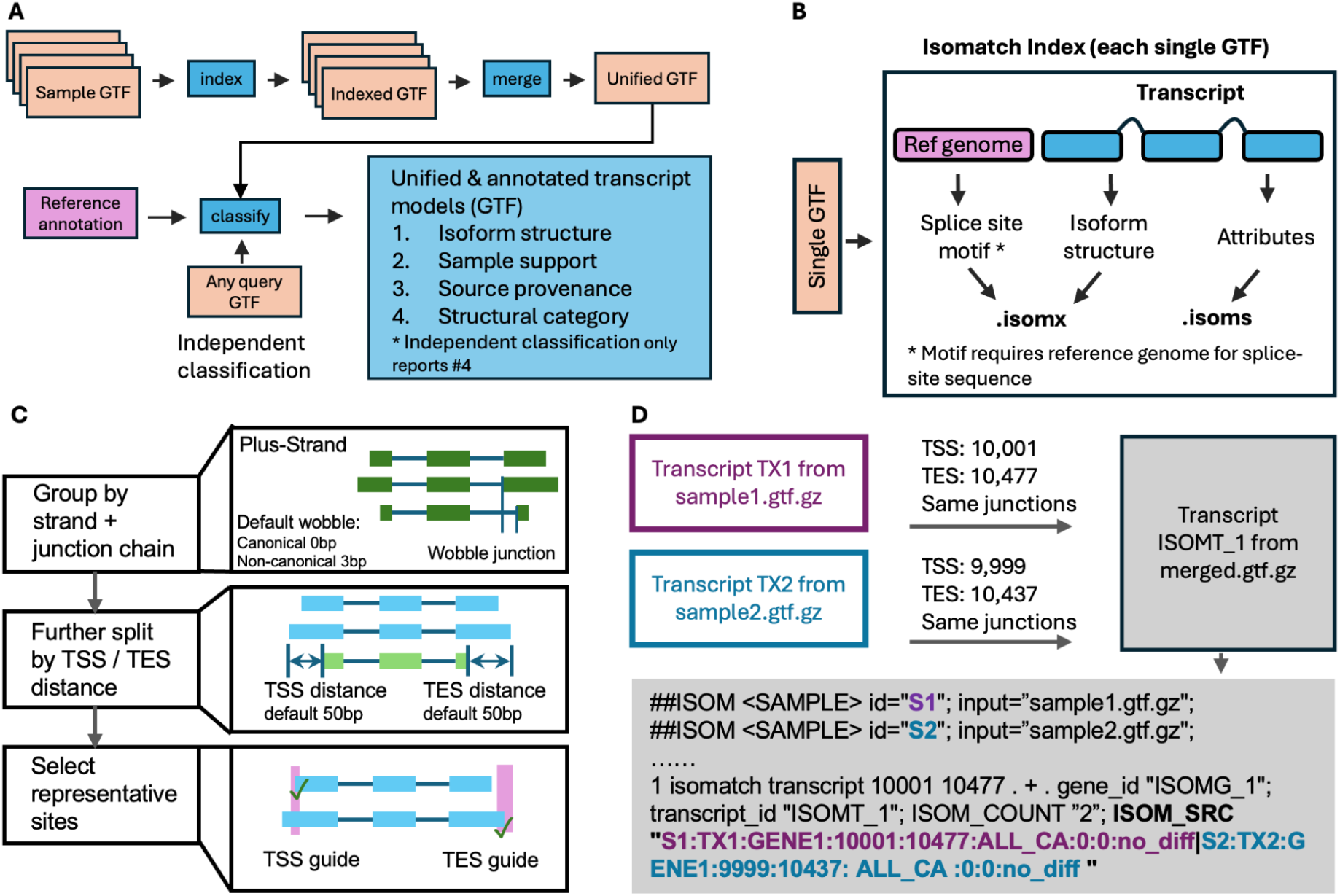
Overview of IsoMatch functionality. **A)** IsoMatch workflow for creating an integrated transcript catalog with classification, population support and tracking information. **B)** Structure of the IsoMatch index. Two files are created per single-sample GTF. The .isomx file stores core isoform structures. The .isoms file stores per-transcript attributes. **C)** Merge strategy implemented in IsoMatch. Transcripts are first grouped by junction-chain identity and strand; a configurable wobble tolerance can be applied at this step to absorb small base-pair offsets at junctions (default: 0 bp for canonical transcripts, 3 bp for non-canonical transcripts). Each group is then further split by TSS and TES distance thresholds (50 bp), and a representative isoform structure is selected (**Methods**). When guided TSS/TES sites are supplied, candidates falling within the guided regions are prioritized. **D)** Encoding of population-scale attributes in standard GTF. Top: two transcripts originating from two single-sample GTFs are merged into a single transcript in the merged GTF. Bottom: an example GTF record generated by IsoMatch. The ##ISOM <SAMPLE> header lines register each source GTF and its sample ID (e.g., S1 for the sample1.gtf.gz). For every merged transcript, the ISOM_SRC attribute records the contributing source transcripts (**Methods**). The format is S{sample}:{tx_id}:{gene_id}:{genome_coordinate_start}:{genome_coordin ate_end}:{tx_type}:{donor_diff_sum}:{acceptor_diff_sum}:{junction_dif fs}.

Beyond transcript merging, IsoMatch preserves transcript-level traceability and connects the merged catalog directly to reference-based structural classification. The resulting merged GTF follows the standard GTF specification, with additional comment-line headers and transcript attributes that provide full traceability directly within the GTF’s attributes column. Among the traceability, the key attribute ISOM_SRC records the source transcript ID and the original input file for each contributing transcript. This record is retained through subsequent classification, enabling any cohort-level transcript model or annotation result to be traced directly to the original GTFs and transcript calls that support it. **Figure 1D** shows an example output in which one merged transcript is supported by two input GTF files. Third, IsoMatch compares each transcript against a specified reference annotation (e.g., GENCODE) and assigns it a SQANTI3-compatible structural category based on its relationship to the reference transcripts. The classify function accepts any GTF file and can therefore be used independently of IsoMatch merging to annotate transcript models generated by other assembly or transcript-discovery workflows. When the input is a GTF produced by IsoMatch merge, the classification output also retains the links between each merged transcript and its source transcripts across samples. Thus, merging and classification are not independent processing steps in IsoMatch, but components of a single traceable workflow built on the same transcript structural representation. Each transcript processed throughout IsoMatch carries a sample support count, a classification relative to the reference annotation, and full tracking back to its source transcripts (**Figure 1A**).

### IsoMatch preserves splice-junction structure during transcript merging

Distinct splice-junction chains capture different patterns of exon connectivity, making their preservation during transcript merging important for characterizing alternative splicing across samples(Jetzinger et al. 2026). We benchmarked IsoMatch and compared it with the state-of-the-art tools for transcript merging and classification. Because no complete ground-truth transcript catalog is available, we used the LRGASP WTC11 benchmark as a proxy. We evaluated transcript structure preservation during merging and concordance in transcript classification across tools(Pardo-Palacios, Wang, et al. 2024). The WTC11 dataset includes six sequencing protocols on the WTC11 cell line, covering the two major long-read sequencing platforms, Pacific Biosciences (PB) and Oxford Nanopore Technologies (ONT), with three technical replicates for each protocol. IsoQuant(Prjibelski et al. 2023) was used to generate transcript sets for each sample with platform-specific parameters, and LRGASP annotation based on GENCODE v38 was used as the reference annotation to guide the transcript discovery(Pardo-Palacios, Wang, et al. 2024). Across all 18 samples, a total of 463,012 transcripts were detected by IsoQuant, with an average of 25,722.89 transcripts per sample. ONT-dRNA detected the fewest transcripts (∼14,612 transcripts per replicate), whereas PacBio-cDNA detected the most (∼36,454.67 transcripts per replicate) (**Figure 2A** and **Table S1**).

**Figure 2.**
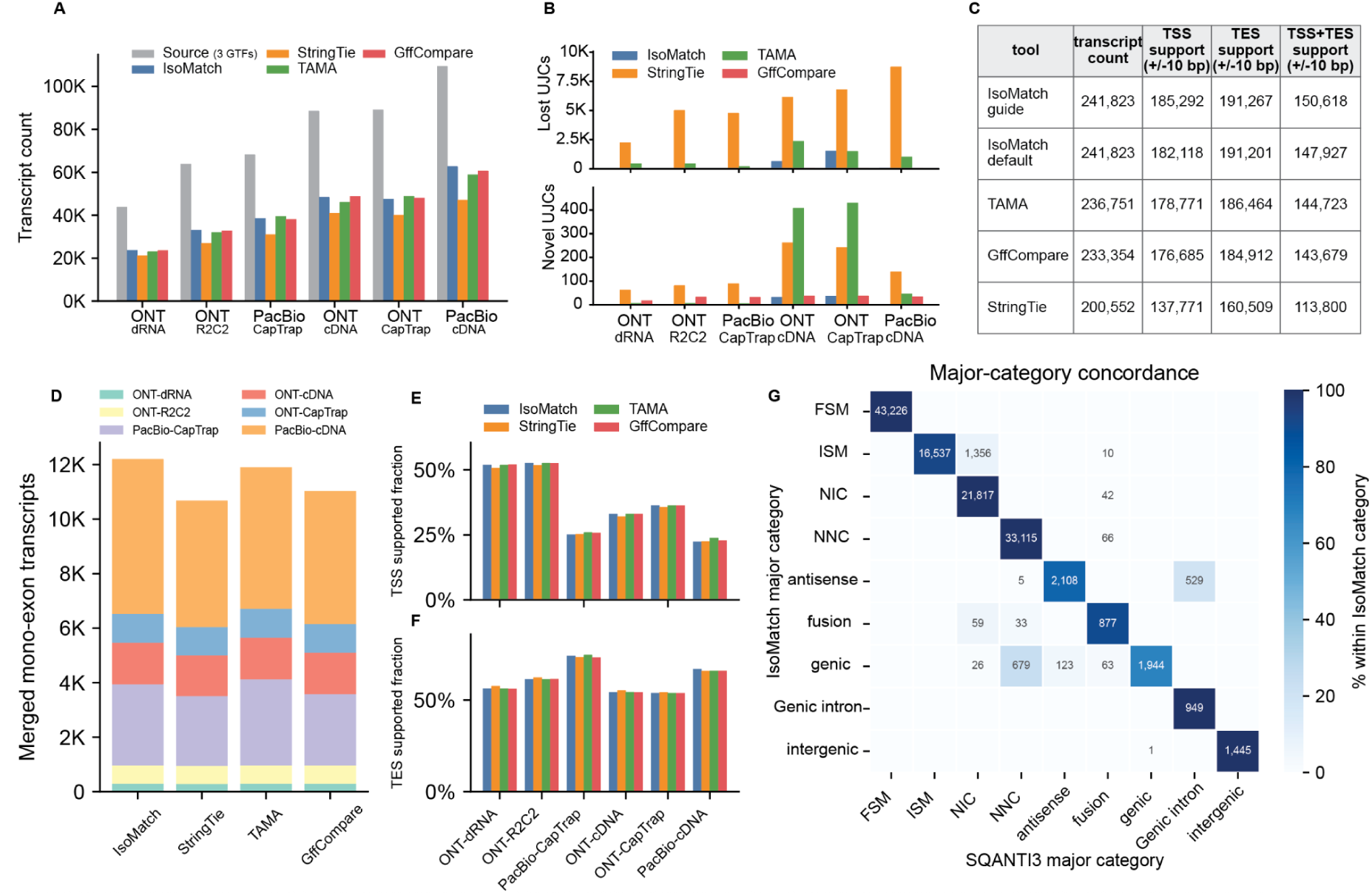
Evaluation of transcript integration and classification using IsoMatch on the LRGASP WTC11 datasets. **A)** Transcript counts before and after merging across six LRGASP WTC11 sequencing protocols. Source transcript models from three technical replicates per protocol were merged using IsoMatch (blue bars), StringTie (yellow bars), TAMA (green bars), and GffCompare (red bars). Grey bars show the total number of transcripts by combining all transcripts in replicates in each group. **B)** Splice-junction-chain changes after merging of WTC11 multi-exon transcripts. For each WTC11 sequencing protocol, three replicate GTFs were filtered to multi-exon transcripts, merged with IsoMatch, StringTie, TAMA, or GffCompare using default parameters, and converted to unique junction chains (UJCs). Bars show input UJCs absent from each merged output (Lost-UJCs, top) and output UJCs not present in the input set (Novel-UJCs, bottom). **C)** Terminal evidence support for WTC11 merged multi-exon transcript ends. The table reports the number of merged multi-exon transcripts from each tool whose TSS, TES overlap external human GRCh38 terminal evidence within a +/- 10 bp in a strand-specific comparison. IsoMatch guide mode uses the same inputs as IsoMatch default with additional TSS/TES guide evidence. The transcript count column reports, for each tool, the total number of multi-exon transcripts across the six protocol-specific merged GTFs. **D)** Number of mono-exon transcripts after merging by each tool. Bars are stacked by WTC11 sequencing protocol, showing each protocol’s contribution to the total merged mono-exon transcript count per tool. **E)** Fraction of merged mono-exon transcripts with TSS support. For each WTC11 protocol, bars compare the four merge tools, counting TSS support within 10 bp of external TSS evidence. **F)** Fraction of merged mono-exon transcripts with TES support. For each WTC11 protocol, bars compare the four merge tools, counting TES support within 10 bp of external TES evidence. **G)** Concordance of major structural classification categories between IsoMatch and SQANTI3 QC using the IsoMatch-merged transcript catalog. The heatmap shows transcript counts and the percentage within each IsoMatch category.

For benchmarking the IsoMatch merge function, we compared IsoMatch with GffCompare(G. Pertea and Pertea 2020), StringTie in the --merge mode(Shinder et al. 2026), and TAMA(Kuo et al. 2020). Overall, with default parameters for all tools, transcript merging reduced the number of transcript models across all protocols and tools by an average of 48.12% (range: 42.13%-57.72%). StringTie consistently produced the largest reductions, whereas the relative ordering of the other tools varied by sequencing protocol (**Figure 2A and Table S1**). These reductions reflect the extent of transcript consolidation but do not, on their own, distinguish the removal of redundant models from the collapse of structurally distinct isoforms.

Next, we divided the transcripts in the 18 WTC11 GTF files into multi-exon and mono-exon categories to obtain further details in the individual merge criteria. This distinction reflects fundamental differences: multi-exon transcripts can be compared based on their splice junction chains, whereas mono-exon transcripts lack splice junctions and are merged primarily based on similarities in terminal boundaries.

For multi-exon transcripts, we asked whether transcript merging altered the set of splice-junction chains present in the input. Preserving this information is important for reliably representing exon connectivity and enabling comparison of individual splice junctions with independent short-read RNA-seq evidence. We represented each distinct splice-junction chain as a unique junction chain (UJC). A merge operation that only consolidates transcripts with identical splice-junction chains should preserve the input UJC set. Deviations from this expectation can occur in two directions: an input UJC can be lost if it is consolidated into a transcript with a different junction chain (lost-UJC), or an output-only UJC can be created if the representative merged transcript contains a junction chain that was not present in any input model (novel-UJC). We therefore quantified both types of change for each merging tool.

Under default parameters, all tools exhibited varying degrees of UJC changes. As shown in **Figure 2B**, GffCompare had the fewest total lost-UJCs across six protocols (257), followed by IsoMatch (2,340), TAMA (6,012), and StringTie (33,744). Further examination of IsoMatch results indicated that all (2,340/2,340) UJCs lost by IsoMatch were attributable to its default splice-site wobble strategy, which absorbs transcripts containing non-canonical splice junctions into nearby canonical transcripts within a 3 bp tolerance. For GffCompare, all 257 lost-UJCs involved removal of an intron shorter than 5 bp, consistent with GffCompare’s close-exon merging behavior. For TAMA, of the 6,012 lost-UJCs, 5,030 (83.67%) involved splice-site differences within TAMA’s 10 bp tolerance, consistent with fuzzy matching. For StringTie, 33,480 of 33,744 lost-UJCs (99.22%) were subsets of longer UJCs, consistent with its shorter-chain absorption strategy. Despite differences in merging strategies across tools, IsoMatch preserves high-quality transcripts with exclusively canonical splice sites while effectively incorporating and consolidating transcripts containing non-canonical splice sites. Notably, IsoMatch can retain all unique junction chains (UJCs) by setting -A 0 -D 0 -U 0. Its default settings are designed to reduce the impact of artifactual transcripts arising from sequencing or alignment errors. Such transcripts often resemble a canonical transcript but differ by 1-3 bp at a small number of splice sites. A detailed analysis of the effects of splice-site wobble parameters is provided in **Supplementary Note 1**.

IsoMatch generated the fewest total novel-UJCs across six protocols (75), followed by GffCompare (196), StringTie (881), and TAMA (907) (**Figure 2B**). Inspection of the IsoMatch output showed that these cases (75/75) arose from representative splice-site selection. When multiple non-canonical transcripts are merged with 3 bp wobble in splice junction, IsoMatch selects the most frequently observed coordinate at each splice site. In a small number of groups, the resulting combination of representative splice sites produced a UJC that was not present as a complete junction chain in any individual input transcript. For GffCompare, all 196 novel-UJCs involved the removal of an intron shorter than 5 bp, consistent with close-exon merging. All 881 StringTie novel-UJCs contained only introns present in the input, consistent with new combinations or subsets of existing intron chains. All TAMA novel-UJCs had a source chain with corresponding splice sites within the default 10 bp tolerance, consistent with fuzzy matching and representative-model selection; a small subset also involved shorter-chain absorption under the capped option (**Methods**).

To determine whether UJC changes reflected the core splice-junction matching behavior or the additional treatment of transcript boundaries, we repeated the benchmark under splice-junction-only settings in which TSS and TES differences were ignored (**Methods**). Under these conditions, both IsoMatch and TAMA preserved the complete input UJC set. For TAMA, achieving junction-only behavior requires highly permissive terminal-boundary parameters because the same boundary settings are applied to both mono- and multi-exon transcripts. We therefore used this configuration only as a controlled test of UJC preservation rather than as a recommended setting for routine transcriptome merging for TAMA.

### IsoMatch retains transcript-boundary diversity during transcript integration

We next asked a different question: whether tools retained multiple transcript models that shared the same UJC but differed at their transcript boundaries. For protein-coding transcripts, differences in TSSs or TESs can alter untranslated regions (UTRs), with potential consequences for mRNA stability, localization and translation(X. Wang et al. 2016; Tushev et al. 2018). GffCompare and StringTie produced a single output transcript for each UJC, whereas IsoMatch and TAMA retained multiple transcripts for 8,759 and 7,291 UJCs, respectively. These UJCs comprised 19,432 transcripts for IsoMatch (mean, 2.22 transcripts per UJC) and 15,732 transcripts for TAMA (mean, 2.16 transcripts per UJC). These models were kept separate because of differences in their TSSs or TESs. Thus, although their splice-junction structures (UJC) were identical, IsoMatch and TAMA preserved terminally distinct transcript models rather than collapsing them solely on the basis of UJC identity.

We next asked whether these retained terminal differences were supported by independent TSS and TES evidence. We performed strand-aware intersections between the TSSs and TESs of merged transcripts and the corresponding evidence intervals in refTSS(Abugessaisa et al. 2019) and PolyASite(Herrmann et al. 2020). Because some evidence intervals span only 1 bp, we used a +/- 10 bp window around each transcript TSS and TES when assessing overlap. We found that 7,366 (84.1%) and 5,885 (80.7%) UJCs retained by IsoMatch and TAMA, respectively, contained at least two transcripts with TSS support from refTSS or TES support from PolyASite. Requiring support for both TSS and TES reduced these numbers to 2,430/8,759 (27.7%) for IsoMatch and 2,127/7,291 (29.2%) for TAMA. These findings provide biological support for retaining terminally distinct isoforms with identical splice junction chains and are consistent with the use of multiple TSSs or TESs by transcripts sharing the same splicing structure(E. T. Wang et al. 2008; Reyes and Huber 2018).

To extend this analysis beyond transcripts sharing a UJC, we evaluated all multi-exon transcripts from the six protocol-specific merges. For each tool, we intersected the TSSs and TESs of the merged transcripts with refTSS and PolyASite regions, respectively, and calculated the proportion supported at both ends across all six protocol-specific outputs. We also evaluated IsoMatch’s guided mode, which accepts external TSS/TES evidence intervals to prioritize supported boundaries when selecting representative transcripts during merging. For each tool, we summed the numbers of transcripts supported at both ends across the six protocol-specific merged catalogs and divided this sum by the total number of output transcripts across those catalogs to calculate the overall support proportion, without deduplicating transcript models across protocols.

Among all evaluated tools, as shown in **Figure 2C**, the two IsoMatch modes produced the highest numbers of transcripts supported at both ends. IsoMatch guide and IsoMatch default yielded 150,618 and 147,927 transcripts supported by both TSS and TES evidence, corresponding to 62.28% and 61.17% of their total output transcripts, respectively. Compared with the default mode, the guide mode produced 2,691 additional transcripts with support at both ends and increased the support rate by 1.11 percentage points. Because the same TSS and TES evidence was used for both guided merging and evaluation, this increase demonstrates effective incorporation of prior information rather than independent validation of transcript boundary accuracy. For comparison, TAMA, GffCompare, and StringTie produced 144,723 (61.13%), 143,679 (61.57%), and 113,800 (56.74%) transcripts with support at both ends, respectively. Overall, IsoMatch guide mode achieved the best performance in terms of both the number and proportion of transcripts with support at both ends, while IsoMatch default mode retained the second-highest number.

We next examined mono-exon transcript merging. Because mono-exon transcripts lack splice junctions, their merging criteria differ substantially from those applied to multi-exon transcripts. IsoMatch merges mono-exon transcripts primarily based on genomic coordinates, strand, and reciprocal overlap (default: 90%). Across the six WTC11 sequencing protocols, IsoMatch retained the largest number of mono-exon transcripts after merging (12,200), followed by TAMA (11,890), GffCompare with the -C option (11,025), and StringTie (10,674). Although mono-exon transcript counts were similar across tools, they varied substantially across sequencing protocols. Mean counts across tools were highest for PacBio-cDNA (5,088.00), followed by PacBio-CapTrap (2,826.25), ONT-cDNA (1,515.75), ONT-CapTrap (1,053.00), ONT-R2C2 (669.75), and ONT-dRNA (294.50) (**Figure 2D**). TSS and TES evidence support among the merged mono-exon transcripts likewise varied more across sequencing protocols than across tools (**Figure 2E** and **2F**). Specifically, ONT-R2C2 and ONT-dRNA showed notably higher proportions of transcripts with TSS support than the other sequencing protocols, whereas PacBio-cDNA and PacBio-CapTrap had the highest proportions with TES support.

Together, these comparisons across multiple WTC11 sequencing protocols and commonly used transcript merging tools indicate that IsoMatch largely preserves splice junction chains during multi-exon transcript merging while consolidating nearby non-canonical junctions through its default splice-site wobble strategy. IsoMatch also retains terminal isoform differences associated with alternative TSSs or TESs, a subset of which are supported by external TSS/TES evidence. For mono-exon transcripts, differences across tools were relatively small, whereas sequencing protocols were associated with larger differences in transcript counts and TSS/TES support.

### IsoMatch achieves high concordance with SQANTI3 while preserving transcript traceability

Finally, we evaluated IsoMatch’s classification module, which compares each query transcript against a specified reference annotation and assigns a SQANTI3-compatible structural category based on its relationship to the reference transcripts. For this benchmark, we used IsoMatch with default settings to merge the 18 WTC11 GTF files into a unified set of 125,010 transcripts as input. We then compared the structural classifications assigned by IsoMatch and SQANTI3 using GENCODE v49 basic as the reference annotation. **Figure 2G** shows that 97.61% of transcripts are concordant for the main structural classification (i.e., assigning the same classification codes) and 99.11% sub-category classification concordance within the main structural category between the two tools(**Table S2**). In particular, all 43,226 transcripts classified as full-splice match (FSM) by IsoMatch were also classified as FSM by SQANTI3. Among the transcripts classified by IsoMatch as incomplete-splice match (ISM), novel-in-catalog (NIC), and novel-not-in-catalog (NNC), 16,537 of 17,903 (92.37%), 21,817 of 21,859 (99.81%), and 33,115 of 33,181 (99.80%), respectively, received the same category from SQANTI3 (**Figure 2G** and **Table S2-S4**).

Among the 2,992 transcripts with discordant major-category assignments, representing 2.39% of the catalog, the differences were concentrated primarily in three category pairs: IsoMatch ISM versus SQANTI3 NIC, IsoMatch genic versus SQANTI3 NNC, and IsoMatch antisense versus SQANTI3 genic intron. Together, these patterns accounted for 85.7% of all discordant assignments (**Figure 2G** and **Table S4**). The ISM category showed 92.37% overlap between the two tools, with 16,537 of 17,903 transcripts classified as ISM by both. Most of the remaining discordance was attributable to 1,356 transcripts classified as ISM by IsoMatch but as NIC by SQANTI3. These transcripts occurred on both strands, and their genomic leftmost terminal exon spanned one or more introns of the reference transcript selected by IsoMatch, while their splice-junction chains formed contiguous subsets of the corresponding reference junction chains. IsoMatch therefore classified them as ISM with intron retention. In contrast, the exon-boundary comparison used by SQANTI3 returned partial rather than subset matches for the same query-reference pairs, and SQANTI3 subsequently assigned these transcripts to NIC, predominantly with the intron-retention subtype (1,335/1,356). This discrepancy reflects differences in how the two tools define ISM when a terminal exon spans reference introns. The other major discordance patterns similarly arose from differences in the evidence and priorities used for gene association and structural classification. Further details are provided in **Supplementary Note 2**.

Together, IsoMatch preserves source-sample and original-transcript identifiers throughout classification, allowing transcripts selected by structural category to be traced back to their contributing input models.

### IsoMatch scales to millions of transcript models with low memory requirements

The rapid growth of transcriptomic datasets, including population-scale long-read studies and large collections of publicly available transcript annotations (e.g. HPRC, ENCODE, 1KGP, and CARD), has created a practical challenge for integrating and annotating transcript models at scale(Lucas et al. 2026; 1000 Genomes Project Consortium et al. 2010; Kouam et al. 2026; Reese et al. 2023). We therefore evaluated the scalability of IsoMatch for large-scale transcript integration and classification using all 163 GTF files downloaded from the ENCODE4 project(Reese et al. 2023). Each file contained an average of 351,565 transcript models, resulting in substantial input size and computational overhead when using conventional plain-text GTF workflows. To improve scalability for large cohorts, IsoMatch natively supports gzip-compressed GTF files, reducing storage and I/O requirements compared with established transcript-merging approaches that require uncompressed input.

We benchmarked the memory usage and CPU time required to merge 10, 20, 50, 100, and 163 GTF files on a workstation with 64 GB of RAM. All tools were run using their default thread settings (1 thread). As shown in **Figure 3A**, only IsoMatch and StringTie successfully merged all 163 GTF files. In contrast, GffCompare encountered an out-of-memory error when merging 100 GTF files, whereas TAMA only completed the merge of 10 input samples. These results highlight the current scalability challenge in the field. Across all comparisons, IsoMatch used substantially less memory than the other tools. For example, when merging 163 GTF files, IsoMatch used 0.49 GB of peak memory, whereas StringTie used 19.12 GB. IsoMatch also showed an advantage in runtime. As shown in **Figure 3B**, IsoMatch required less CPU time than the other tools across all tested input sizes. In particular, when merging 163 GTF files, IsoMatch took 959.15 seconds, while StringTie took 1,944.4 seconds. Given the low computational resource usage observed when merging 163 GTF files, we further tested the scalability of IsoMatch using a synthetic duplicated-input stress test. For this benchmark, we used 1,000 identical GTF files (GENCODE v49 basic GTF, containing 280,000 transcripts) as input. IsoMatch completed the merge in 17 h 53 min with only 5.24 GB of peak memory, whereas StringTie failed to load all 1,000 inputs. Thus, the scalability advantage is not limited to reduced runtime or memory usage, but expands the size of transcript collections that can be practically integrated within a single analysis.

**Figure 3.**
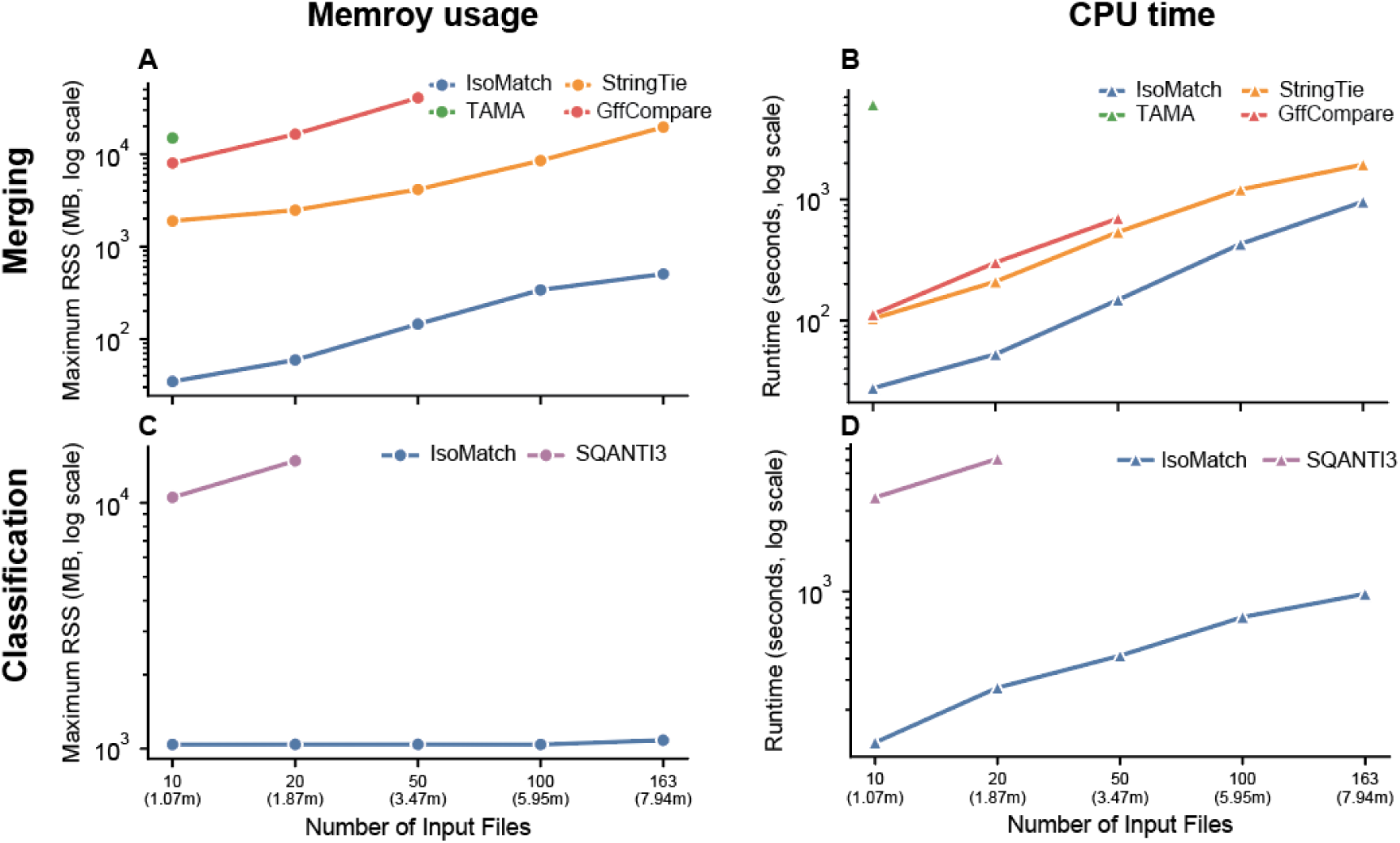
Scalability of IsoMatch for population-scale transcript integration and classification. **(A, B)** Peak memory usage and CPU runtime for merging 10, 20, 50, 100, and 163 ENCODE GTF files using IsoMatch, StringTie, TAMA, and GffCompare on a workstation with 64 GB RAM. The x-axis indicates the number of input GTF files; labels beneath panels C and D additionally give the number of transcripts in the corresponding IsoMatch-merged catalogs. **(C, D)** Peak memory usage and CPU runtime for classifying the corresponding merged GTF files using IsoMatch and SQANTI3. Memory usage and runtime are shown on log-scaled axes. IsoMatch completed all merge and classification benchmarks with lower computational resource usage, whereas other tools (except StringTie) failed on larger input sets. Missing points indicate runs that did not complete. The numbers in parentheses indicate the number of transcripts, in millions. The x-axis indicates the number of input GTF files; labels beneath panels C and D additionally give the number of transcripts in the corresponding IsoMatch-merged catalogs. RSS (Resident Set Size): The physical RAM occupied by a process.

We further evaluated the classification function of IsoMatch and compared it with SQANTI3. SQANTI3 QC performs additional quality-control analyses beyond structural classification. This comparison therefore measures the computational resources required to obtain structural classifications using the evaluated workflows, rather than the efficiency of equivalent classification-only tasks. Specifically, we classified the five GTF files containing 1,066,077-7,938,545 transcripts generated in the previous step by IsoMatch and recorded the peak memory usage and CPU runtime for each task. As shown in **Figures 3C** and **3D**, SQANTI3 successfully completed the classification of only the GTF files generated from the merged 10-sample and 20-sample datasets. In contrast, IsoMatch successfully classified all merged GTF files. Notably, IsoMatch required lower memory usage and shorter runtime across all comparable GTF files, including the 10-sample merged GTF (1.02 GB, 127.8 seconds) and the 20-sample merged GTF (1.02 GB, 269.73 seconds) (**Table S5**). For the largest classification task, involving the GTF merged from 163 samples, IsoMatch completed the analysis with a peak memory usage of 1.06 GB and a runtime of 967.84 seconds.

In summary, these results demonstrate that IsoMatch is well suited for future large-scale transcriptome studies, combining modest computational requirements with the ability to process 1,000 GTF inputs in a synthetic stress test, while providing a consistent framework for isoform merging and annotation that preserves full traceability from source transcripts to their final merged and classified representations.

### Reference transcript annotations remain substantially discordant under multiple equivalence criteria

Few studies have systematically examined the differences among reference annotations and their impact on transcript classification. Thus utilizing IsoMatch, we integrated three commonly used reference transcript catalogs, including GENCODE v49 (basic), RefSeq v110 (release 110), and CHESS v3.1.3(Varabyou et al. 2023; Mudge et al. 2025; Goldfarb et al. 2025). As shown in **Figure 4A**, GENCODE v49 basic contains 280,000 transcripts, RefSeq v110 contains 179,144 transcripts and CHESS v3.1.3 contains 158,066 transcripts. After merging with IsoMatch using default parameters (0-bp splice-site tolerance for canonical transcripts, 3-bp tolerance for transcripts containing non-canonical junctions, 50-bp TSS/TES tolerances, and ≥90% reciprocal overlap for mono-exon transcripts), we obtained 464,617 merged transcripts, corresponding to a reduction rate of 24.72% (**Figure 4A**). We observed substantial discordance among the three reference catalogs. Only 35,979 transcripts were shared by all three catalogs, accounting for 7.74% of the merged transcript set, while 79,793 transcripts were shared by two catalogs, accounting for 17.17%. Transcripts unique to a single reference catalog accounted for 49.36%, 16.04% and 9.69% of the merged catalog for GENCODE, RefSeq, and CHESS, respectively (**Figure 4A**). Such substantial discordance among reference annotations has also been independently reported in previous studies(Morales et al. 2022; M. Pertea et al. 2018; Hamaguchi et al. 2021).

**Figure 4.**
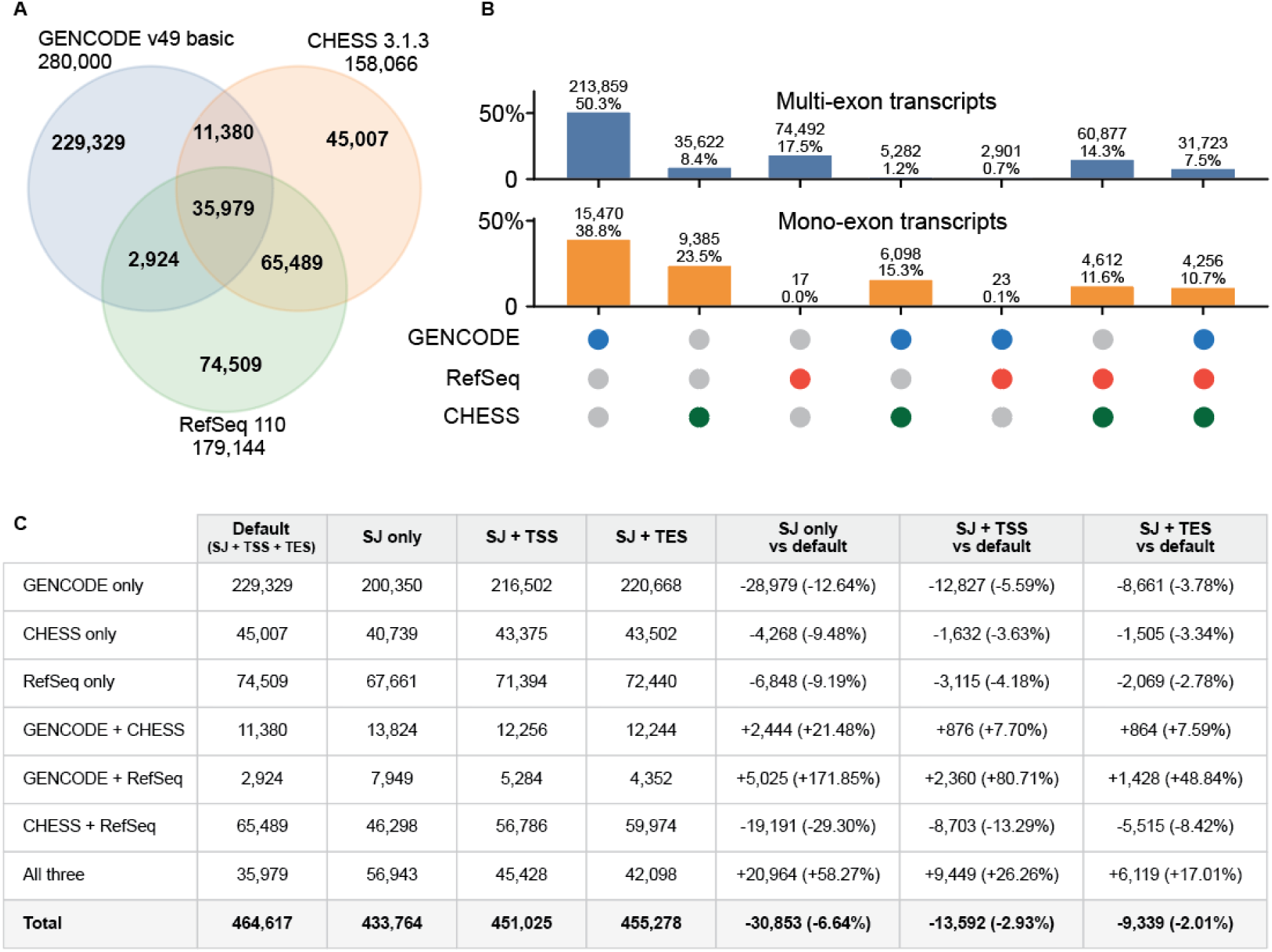
Comparison of reference annotation with IsoMatch. **A)** Overlap among GENCODE, CHESS, and RefSeq transcript annotations. Numbers under each annotation label indicate input transcript records; numbers within each sector indicate unique or shared merged transcript groups, which may consolidate multiple records from the same annotation. **B)** Comparison of annotation overlap patterns between multi-exon and mono-exon transcripts. Bars show the percentage of transcripts assigned to each annotation-sharing category, with the corresponding transcript counts and percentages indicated above each bar. **C)** Effect of transcript-matching criteria (e.g., SJ only) on the number of merged transcripts assigned to each annotation-sharing category (e.g., GENCODE only). Results obtained using default (SJ + TSS + TES), splice junctions alone (SJ only), splice junctions together with transcription start sites (SJ + TSS), or splice junctions together with transcription end sites (SJ + TES). The three rightmost columns show the absolute and percentage changes in transcript counts within each annotation-sharing category under the corresponding transcript-matching criteria relative to the default criteria. Negative values indicate fewer transcripts in that category than under the default criteria, whereas positive values indicate more transcripts.

To further evaluate the robustness of IsoMatch in merging well-established cross-database transcript models, we specifically examined “Matched Annotation from NCBI and EMBL-EBI” (MANE select) transcripts shared between GENCODE and RefSeq(Morales et al. 2022). MANE select transcripts are curated representative transcript models with particular importance for clinical genomics and biomedical research. In GENCODE v49, 19,342 transcripts are annotated as MANE select transcripts, whereas 19,060 MANE select transcripts are present in RefSeq v110. Using the MANE cross-reference (db_xref attribute) provided in the RefSeq GTF, we identified 18,624 MANE select transcripts that could be directly linked between GENCODE and RefSeq. IsoMatch assigned the corresponding GENCODE and RefSeq records to the same merged group for all 18,624 directly linked MANE select pairs, suggesting IsoMatch correctly recovers these known cross-database equivalences. These groups also contained 278 additional non-MANE select transcripts, including 196 from GENCODE and 82 from RefSeq. Of these 278 transcripts, 112 (40.29%) were structurally identical to the corresponding MANE select transcripts, 99 (35.61%) shared the same junction chain but differed by <= 50 bp at transcript termini, 58 (20.86%) were incorporated through splice-junction wobble involving non-canonical splice sites, and nine (3.24%) were mono-exon transcripts with ≥ 90% reciprocal overlap. Thus, all directly linked MANE select pairs were consolidated, while additional group members reflected structural redundancy or differences accommodated by the configured matching criteria.

We next quantified transcripts unique to each of the three reference annotations or shared among them after merging. Briefly, we parsed the ISOM_SRC tracking attribute from the merged GTF to determine whether each merged transcript was supported by GENCODE, CHESS, RefSeq, or combinations thereof. We stratified transcripts by exon number and separately examined the overlap patterns of multi-exon and mono-exon transcripts. In total, the integrated annotation contained 424,756 multi-exon transcripts and 39,861 mono-exon transcripts, which showed markedly different patterns of annotation overlap (**Figure 4B**). GENCODE-only transcripts represented the largest category in both groups, accounting for 50.3% of multi-exon transcripts and 38.8% of mono-exon transcripts. In contrast, CHESS- and RefSeq-specific transcripts showed distinct distributions: CHESS-only transcripts were substantially enriched among mono-exon transcripts (23.5% versus 8.4% for multi-exon transcripts), whereas RefSeq-only transcripts were almost exclusively multi-exon (17.5% versus <0.1% for mono-exon transcripts). Similar differences were observed among transcripts shared by two annotations. Transcripts shared between GENCODE and CHESS but absent from RefSeq accounted for 15.3% of mono-exon transcripts but only 1.2% of multi-exon transcripts. Conversely, transcripts shared between GENCODE and RefSeq but absent from CHESS were rare in both groups, particularly among mono-exon transcripts (0.7% of multi-exon transcripts and 0.1% of mono-exon transcripts). Transcripts shared between CHESS and RefSeq were more evenly represented, comprising 14.3% of multi-exon and 11.6% of mono-exon transcripts. Finally, 31,723 multi-exon transcripts and 4,256 mono-exon transcripts were shared across all three reference annotations, corresponding to 7.5% and 10.7% of the respective groups.

We further evaluated how terminal constraints affect reference transcript merging (**Figure 4C**). When only splice-junction structure was considered, the number of merged transcript groups decreased from 464,617 to 433,764, a reduction of 30,853 groups (-6.64%) relative to the default mode (considers splice junction and TSS/TES). Retaining either TSS or TES partially restored the default grouping, with SJ + TSS and SJ + TES producing 451,025 (-2.93%) and 455,278 (-2.01%) groups, respectively. These results indicate that TSS/TES differences account for a measurable fraction of transcript group separation beyond splice-junction structure, with TES showing a slightly stronger contribution than TSS in this comparison. Relaxing terminal constraints also shifted transcript groups toward broader cross-reference support: the all-three overlap category increased by 20,964 groups (+58.27%) in the SJ-only mode, while single-reference categories decreased. This suggests that a certain fraction of transcripts across GENCODE, RefSeq, and CHESS share splice-junction structures but differ in annotated transcript terminals. However, even under the SJ-only mode, 308,750 merged transcript groups (71.18%) remained specific to a single reference annotation, indicating that splice-junction differences still represent the dominant source of divergence among the three reference transcriptomes.

Together, these results highlight the need for a unified reference annotation that can reconcile both shared and reference-specific transcript structures.

### Transcript novelty depends on reference annotation choice

Given that GENCODE, RefSeq, and CHESS each provide high-quality transcript models but differ substantially in transcript composition, we generated a unified reference annotation (URA) by merging the three catalogs. We expect that this URA provides a more comprehensive reference transcript set for query transcript classification while retaining links to the source annotations through IsoMatch’s transcript tracking functionality. Importantly, clinically relevant transcripts, including MANE select transcripts, were correctly consolidated across annotations, supporting the reliability of IsoMatch for integrating heterogeneous reference transcript catalogs.

To compare classification using individual reference annotations and the URA, we used each catalog as reference annotation to classify transcript models generated from the 119 PacBio long-read RNA-seq datasets from the 1000 Genomes Project(1000 Genomes Project Consortium et al. 2015). To avoid bias toward any reference annotation, we used StringTie in unguided mode to construct reference-annotation-agnostic transcript sets for each sample (**Methods**). As a result, StringTie identified a total of 11,140,448 transcript models, with an average of 93,617 transcripts per sample (range 70,800-137,929). IsoMatch merged these 119 sample-level GTF files in 2 min 10 s (excluding indexing, which is reusable and can be parallelized), yielding a population transcript catalog containing 4,839,988 nonredundant transcript models with default configuration. This 56.6% reduction indicates that many transcript models recur across individuals and can therefore be represented by shared population-level structures.

We next used the sample-level traceability retained by IsoMatch to demonstrate a direct downstream benefit of traceable merging: population support for every merged transcript could be calculated without reconstructing transcript-to-sample relationships directly. The population catalog exhibited a strongly skewed frequency distribution, with most transcripts detected in only one or a small number of samples and a much smaller subset recurrent across many individuals, as shown in **Figure 5A**. Among the 4,839,988 transcripts in the 119-sample catalog, 3,697,027 (76.39%) were detected in a single sample, whereas 1,142,961 were recurrently detected in two or more samples. Furthermore, 155,482 (3.21%) transcripts were supported by at least 10 samples, and 15,344 (0.32%) were detected in more than half of the cohort, corresponding to at least 60 samples. It is worth noting that these transcript models were assembled without reference annotation guidance, which may partly explain the relatively low recurrence observed across samples.

**Figure 5.**
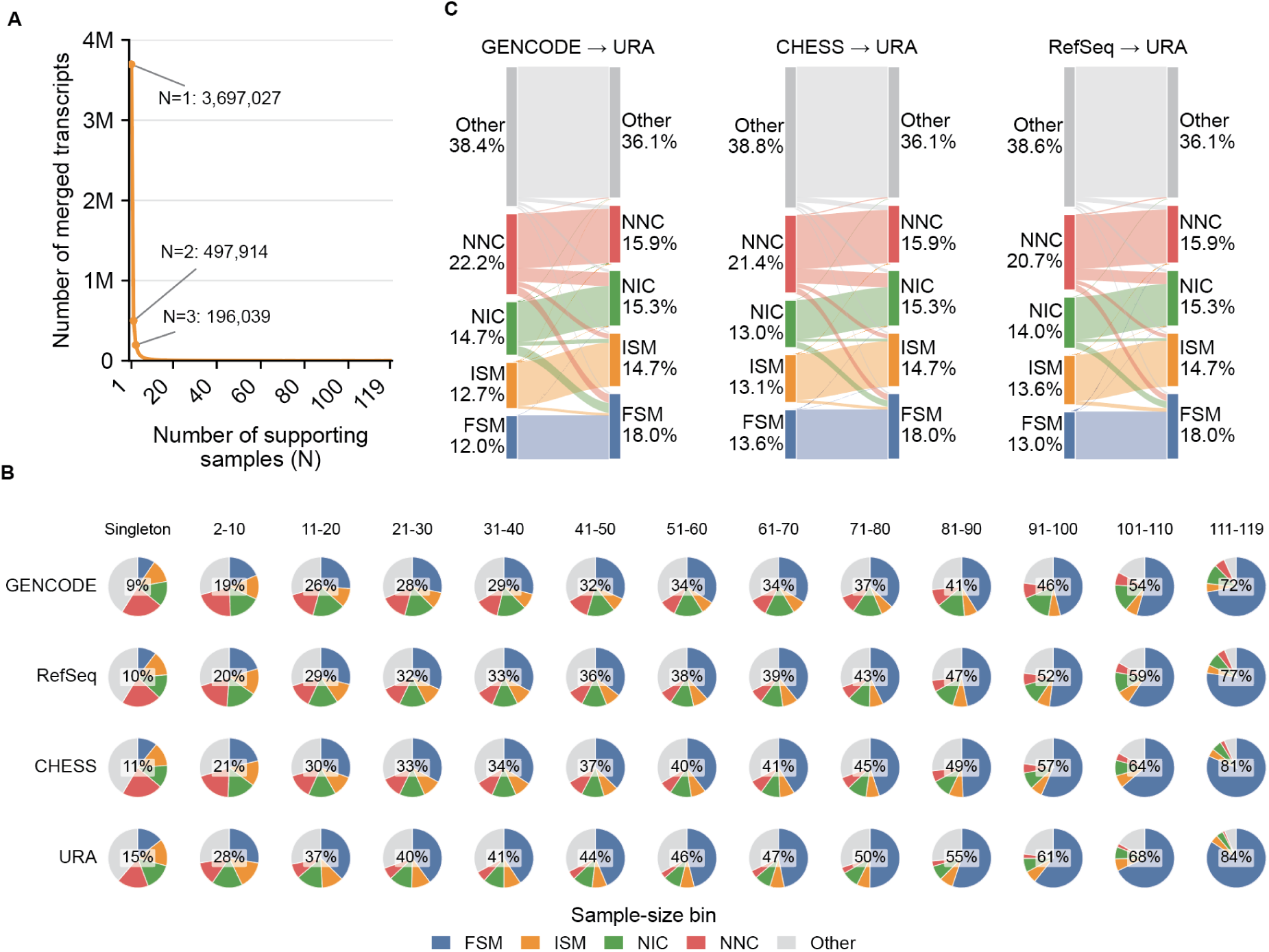
Unified reference annotation facilitates the interpretation of novel isoforms. **A)** Distribution of sample support for merged transcripts across 119 unguided transcriptome assemblies from the 1000 Genomes Project. The x-axis shows the number of supporting samples (N) in ascending order, and the y-axis shows the number of merged transcripts supported by exactly N samples on a linear scale, in millions (M). Points at N = 1, 2, and 3 are annotated with their transcript counts. **B)** Relationship between transcript population recurrence and structural classification. Unified transcripts from 119 samples were grouped by the number of supporting samples, from singletons to transcripts detected in 111-119 samples, and independently classified against GENCODE, RefSeq, CHESS, and the URA. Each pie chart shows the proportion of transcripts classified as full-splice match (FSM), incomplete-splice match (ISM), novel-in-catalog (NIC), novel-not-in-catalog (NNC), or other classes within each sample-size bin. Percentages shown inside the pies indicate the proportion of FSM transcripts. **C)** Sankey plot tracking changes in transcript structural classification when the same transcripts are annotated using individual reference annotations or the URA. “Other” includes classification categories other than FSM, ISM, NIC, and NNC.

We next examined the relationship between population recurrence and structural classification using each individual reference annotation and the URA. Transcripts observed across more samples were increasingly likely to match existing annotations. Among 2,066 transcripts detected in 111-119 samples, 72-84% were classified as FSM, compared with approximately 19-28% among transcripts detected in 2-10 samples and 9-15% among singletons (**Figure 5B, Table S6**). Nevertheless, NIC and NNC transcripts remained among recurrent transcript models, indicating that population recurrence and reference classification capture complementary properties of the merged transcript catalog. Importantly, the population-level transcript models remain directly traceable to their contributing samples and original transcript calls. IsoMatch retains this information within the merged GTF through the ISOM_SRC attribute, linking each unified transcript to its source files and transcript identifiers. Thus, any recurrent, reference-discordant or otherwise selected transcript can be traced directly back to the samples and original models supporting it. This enables subsequent cross-checking across samples, cohorts or analysis workflows and allows population-level observations to be linked back to the underlying transcript calls without relying on separate tracking files.

Finally, the URA consistently increased reference concordance across population-frequency groups compared with any individual annotation (**Figure 5B**). Across the complete population catalog, FSMs accounted for 12.0-13.6% (average 12.9%) of transcripts when classified against individual annotations, compared with 18.0% using the URA (**Figure 5C**). The additional FSM assignments were primarily transcripts classified as ISM, NIC or NNC against individual references. In summary, these analyses demonstrate that IsoMatch can integrate population support, reference classification and source tracing within a single traceable transcript catalog.

## Discussion

RNA sequencing studies increasingly generate transcript models across hundreds to thousands of samples, while the growing availability of public datasets creates new opportunities to compare transcript structures across studies(GTEx Consortium 2013; Reese et al. 2023). This creates a practical challenge that extends beyond transcript discovery within individual samples. Large collections of transcript models need to be integrated at scale, compared against reference annotations using consistent structural criteria and, importantly, remain traceable to the samples and original transcript identifiers from which they originated. These requirements are independent of the sequencing technology or transcript discovery method used to generate the models. IsoMatch addresses these requirements directly at the GTF level, providing a scalable and traceable framework that links transcript merging and reference-based classification within a consistent framework. The benchmarks presented here use long-read-derived transcript models, but IsoMatch itself operates on transcript structures encoded in GTF and does not depend on the sequencing technology or transcript-discovery method that generated them. Transcript catalogs produced by short-read assembly, long-read sequencing, hybrid approaches, annotation pipelines or independent studies can therefore be integrated and classified within the same framework.

A central challenge in such analyses is determining when transcript models should be considered equivalent. Overly permissive merging can collapse distinct isoforms and remove genuine transcript diversity, whereas overly stringent matching can separate equivalent structures because of small differences in splice junctions or transcript boundaries. Our benchmarking suggests that IsoMatch achieves a favorable balance between these two failure modes. In the WTC11 proxy benchmark, IsoMatch preserved splice-junction structure with high concordance between the input and merged unique junction chain (UJC) sets, with the few changes observed attributable to its default non-canonical wobble strategy rather than to indiscriminate consolidation. At the same time, IsoMatch retained transcript models that shared an identical UJC but differed at their transcript boundaries, and a substantial proportion of these terminal differences were supported by independent TSS and TES evidence, indicating that they reflect genuine isoform diversity rather than merging noise. Multi-exon and mono-exon transcripts require substantially different merging criteria, and for mono-exon transcripts we observed comparable performance across tools but pronounced differences across sequencing protocols, suggesting that mono-exon merging is constrained more by the upstream library preparation than by the merging strategy itself. Nevertheless, benchmarking transcript merging remains challenging because no complete ground-truth human transcript catalog exists. Our analyses therefore evaluate identifiable merging inconsistency rather than absolute transcript accuracy, and highlight the need for future benchmarks containing known transcript mixtures and experimentally validated structures.

Our results further quantitatively demonstrate that reference annotation itself substantially influences transcript classification. GENCODE, RefSeq and CHESS contain overlapping but distinct transcript sets, and substantial differences remain even when transcript terminal constraints are relaxed. Consequently, the same experimentally observed transcript can receive different classifications depending on the reference annotation used. Integrating the three annotations with IsoMatch increased the fraction of population transcripts classified as full-splice matches from approximately 12.9% to 18.0%, with transcripts previously classified as ISM, NIC or NNC becoming reference-supported. The resulting unified reference annotation is not intended to define a definitive human transcriptome. Rather, it demonstrates that integrating complementary annotations can reduce classification differences caused simply by reference choice while retaining information about which annotation sources support each transcript model.

Traceability is not merely an output convenience. Once transcript models from multiple samples or catalogs are merged into a cohort-level catalog, downstream filtering, recurrence analysis, reclassification, validation and locus-specific follow-up depend on knowing which original observations support each merged model. An important feature of IsoMatch is that transcript integration does not remove the connection to the underlying data. Each merged transcript retains its source samples and original transcript identifiers directly within the GTF, allowing population-level models to be traced back to their contributing observations. This enables subsequent analyses to ask not only whether a transcript is present in a merged catalog, but where it was observed and which original models contributed to it. In the 1000 Genomes Project analysis, this information allowed us to quantify transcript recurrence across individuals and showed that transcripts observed across more samples were increasingly likely to match existing annotations, while recurrent NIC and NNC structures remained after classification against the integrated reference. More generally, preserving tracing information enables cross-checking of transcript structures across samples, cohorts or analysis workflows and provides a direct link between merged transcript catalogs and downstream sample-level analyses. This becomes particularly important when catalogs are subsequently filtered, reclassified or compared across studies, as the evidence underlying any merged transcript can be recovered without maintaining separate tracking files.

The declining cost of RNA sequencing, expanding cohort sizes, and continued accumulation of public transcriptomic datasets place increasing demands on computational scalability. Comparisons that once involved a handful of transcriptomes may increasingly require integration of hundreds or thousands of GTF files across samples, tissues or independent studies. In our benchmarks, IsoMatch processed large transcript collections with substantially lower memory usage and shorter runtimes than existing tools and completed analyses that other methods could not finish within the available computational resources. Together with configurable transcript matching, reference-based classification and transcript traceability, this scalability enables IsoMatch to provide a general framework for integrating transcript models across modern RNA sequencing studies. Importantly, this scalability extends traceable transcript integration to large cohorts, making IsoMatch applicable across a broader range of study sizes and designs while preserving links to the source transcripts.

Together, IsoMatch makes large-scale transcript models integration computationally feasible, preserves the traceability required for downstream analysis, and provides a consistent framework linking transcript merging with reference-based classification. These capabilities provide a practical foundation for interpreting transcript diversity and reference-dependent novelty at population scale.

## Methods

### 1. IsoMatch workflow

IsoMatch is a standalone command-line tool implemented in Rust, with three main subcommands: index, merge, and classify. Unlike tools that load all transcript records into memory, IsoMatch uses an index-based architecture to reduce memory requirements, supporting large-scale transcript merging and classification. The index subcommand builds a transcript structure index for each GTF file, enabling the merge and classify subcommands to load only the transcripts needed for processing. The merge and classify share the same structural primitives (strand, junction chain, TSS, and TES), but each uses explicit, configurable matching criteria (see below for details). A typical IsoMatch workflow first builds indexes, then merges GTFs into a unified nonredundant transcript set, and finally classifies that set against a reference annotation. This unified workflow bridges transcript integration and classification within a consistent workflow, while enabling rapid, fine-grained parameter tuning to optimize results even for large datasets.

### 2. Structural matching criteria

For merging, IsoMatch assesses transcript equivalence by comparing strand, splice junction chains, donor and acceptor coordinates, and TSS and TES coordinates. Configurable tolerances allow positional differences at splice sites and transcript ends. For each splice junction, donor and acceptor coordinates are compared independently, with separate parameters controlling their respective tolerances. Both canonical splice-site tolerances default to 0 bp, requiring identical splice-junction chains for equivalence. For non-canonical splice sites, IsoMatch allows a default wobble of 3 bp. TSS and TES distance thresholds both default to 50 bp and are used to further separate transcripts into distinct groups.

For classification, IsoMatch uses exact splice-junction matching, consistent with SQANTI3 QC. A query junction matches a reference junction only when both donor and acceptor coordinates are identical. The relationship between the query and reference splice junction chains is used to determine the main structural category. TSS and TES tolerances default to 50 bp and are used only to distinguish subcategories, such as reference_match versus alternative 5′ or 3′ ends.

IsoMatch allows splice-junction wobble during merging to accommodate sequencing and alignment noise across samples. In the standard workflow, this variability is resolved through representative structure selection, making additional tolerance during classification redundant. Classification therefore treats remaining junction differences between the query and a curated reference as structural differences rather than noise, using exact matching to ensure direct comparability with SQANTI3.

### 3. IsoMatch index

IsoMatch builds an index for each input GTF file to enable rapid sequential access with minimal memory requirements. When merge or classify is executed on a GTF that has not been indexed, the index is constructed automatically. The index is reusable and can be constructed individually for each sample. When constructing the index, transcript structures are parsed and sorted by chromosome name and leftmost genomic coordinate. The IsoMatch indexing step stores transcript structures in an isomx file and associated metadata, including transcript attributes, in a separate isoms file. Record offsets link each transcript in the isomx file to its metadata, enabling on-demand retrieval and reducing memory usage during merging and classification. Indexing also requires the corresponding reference genome to classify each splice junction as canonical (GT-AG, GC-AG, or AT-AC) or non-canonical. During merging, these labels distinguish transcripts containing only canonical splice junctions from those containing at least one non-canonical junction, allowing separate merging parameters to be applied to each group. This sequence-context-aware design enables merging criteria to be tailored to splice-junction canonicality as an indicator of transcript confidence.

After indexing, IsoMatch outputs a statistics file summarizing transcript counts overall and by strand, mono- and multi-exon transcript counts, and the proportions of canonical transcripts and splice sites.

### 4. IsoMatch merge function

IsoMatch merge takes multiple indexed GTF files as input and produces a unified, nonredundant transcript set in standard GTF format, with specialized attributes that preserve source-transcript traceability.

IsoMatch processes transcripts chromosome by chromosome. Within each chromosome, it uses a k-way merge algorithm to identify initial groups of transcripts that overlap by at least 1 bp. Each group is then divided into subgroups by strand and number of splice junctions. Within each subgroup, transcripts are further partitioned by splice-junction canonicality. Transcripts containing only canonical junctions are merged first to form a backbone into which transcripts containing one or more non-canonical junctions can subsequently be absorbed.

All-canonical transcripts are grouped by comparing their splice junction chains, TSSs, and TESs using configurable tolerances for donor sites (-d), acceptor sites (-a), unstranded transcripts (-u), TSSs (-s), and TESs (-e). Transcripts containing non-canonical junctions are then matched to the resulting all-canonical groups using a separate set of tolerances (-D, -A, -U, -S, and -E). Any remaining transcripts that cannot be absorbed are grouped separately using the same parameters.

After the source transcripts have been grouped, IsoMatch constructs a representative transcript structure for each group. Because source transcripts may differ slightly in their coordinates, IsoMatch supports several policies for selecting representative sites. For splice junctions, the available policies are major (the most frequent coordinate; default), longer (the coordinate defining the longer exon), and shorter (the coordinate defining the shorter exon). When reference TSS or TES annotations, such as refTSS or PolyASite, are provided, the transcript ends closest to known sites are preferentially selected. Each representative transcript will be assigned a transcript ID.

Mono-exon transcripts are processed separately based on strand and reciprocal overlap, with the overlap threshold controlled by --mono-ovlp.

#### Identification of gene clusters

IsoMatch identifies the gene cluster after all source transcripts have been merged into representative transcripts. Since the transcripts from the last step k-way merge may come from different genes, for each genomic super-cluster, merged transcript groups are first ordered by strand and genomic span. Transcripts on the same strand are partitioned into overlapping span groups, then further connected by shared splice sites. Splice-connected groups are treated as candidate genes, with an additional heuristic to separate potential read-through structures when a clear low-coverage valley exists between two transcript-dense regions.

Transcripts that cannot be assigned through splice-site connectivity, such as mono-exon or splice-isolated models, are assigned by exon overlap. They inherit an existing gene group only when they overlap it above a minimum threshold and the best match is unambiguous. Remaining unassigned transcripts are grouped together using reciprocal exon-overlap criteria. Each resulting gene group receives a new sequential merged gene identifier.

#### Output of IsoMatch merge

IsoMatch outputs merged transcripts in a GTF file that is compatible with the GTF specification. In addition, IsoMatch adds custom header lines and transcript-level attributes to record metadata for each merged transcript. These metadata fall into two main categories: source-transcript traceability and summary statistics.

For traceability tracking, IsoMatch writes header lines that register each input GTF file and assigns it a sample label beginning with S, such as S1 or S2. For each merged transcript, ISOM_SRC records the traceability of every source transcript merged into an output transcript: one record per source, joined by |, each a colon-separated block of nine fields: S{sample}:{tx_id}:{gene_id}:{genome_coordinate_start}:{genome_coordinate_end}:{tx_type}:{donor_diff_sum}:{acceptor_diff_sum}:{junction_d iffs}. S{sample} is the 1-based input-file index from the ##ISOM <SAMPLE> headers; tx_id, gene_id, start and end are taken verbatim from the source GTF (1-based, inclusive); tx_type is the splice-site class (MONO, ALL_CA, PRT_CA, NOT_CA). MONO denotes mono-exon transcripts; ALL_CA, PRT_CA, and NOT_CA denote transcripts with canonical splice motifs at all, some, or none of their junctions, respectively. donor_diff_sum and acceptor_diff_sum are the summed absolute distances, in bp, between the source and representative splice sites on each side, assigned strand-aware (the donor is the left junction coordinate on the plus strand, the right one on the minus strand). junction_diffs lists only the junctions that differ, as 1-based (junction_number,left_offset,right_offset) tuples, or no_diff; offsets are representative - source, so the source coordinate can be recovered.

The second category of metadata summarizes properties of the merged transcript. IsoMatch records the number of exons (ISOM_EXONS), the number of source transcripts merged into the output transcript (ISOM_COUNT), the number and fraction of input samples represented by the transcript (ISOM_SAMPLE_CNT and ISOM_SAMPLE_FREQ), and the representative-selection strategy used for splice junctions, TSS, and TES (ISOM_REPR_POLICY). Available policies are major (the most frequent coordinate; default), longer (the coordinate defining the longer exon), and shorter (the coordinate defining the shorter exon). These attributes make the merged GTF both annotation-compatible and directly interpretable for downstream analysis.

#### Evidence files for TSS and TES

IsoMatch provides preprocessed evidence files for human transcription start sites (TSSs) and transcript end sites (TESs). TSS annotations from refTSS v4.1 were downloaded and converted to BED format with six columns: chromosome, start, end, ID, score, and strand. TES annotations were downloaded from the PolyASite database and converted to the same BED format. The processed evidence files are available in the evidence directory of the IsoMatch repository.

### 5. IsoMatch classify function

IsoMatch classify takes a query GTF and the reference annotation (GTF) as input and outputs a classification table along with an annotated GTF. The merged GTF from the IsoMatch merge can be seamlessly used as input but it can also accept any other GTF files. However the query GTF and the reference annotation GTF must use the same reference genome. For each query transcript, IsoMatch will find all overlapped reference transcripts and compare the query transcript with all overlapped reference transcripts one by one. If multiple reference transcripts are found, the best hit will be selected and written to output. The detailed steps are listed below.

The reference annotation is parsed once into a set of per-chromosome catalogs. For each chromosome, all splice junction pairs and donors and acceptors are collected from all transcripts. Additionally, for each gene (defined by gene_id in the reference annotation GTF), splice-junction pairs and TSS/TES coordinates are collected and deduplicated. Those coordinates represent a non-redundant catalog that will be used in next steps to evaluate each query transcript and assign the corresponding evidence to them.

Before conducting query-to-reference pairwise comparisons, IsoMatch will evaluate each query transcript with the catalogs built from the last step. Briefly, IsoMatch assesses 1) whether all donor and acceptor sites in the query are present anywhere in the annotation, regardless of their original transcript or gene, 2) whether any query exon spans a known intron, indicating intron retention, 3) whether the entire query lies within a known intron. These annotation-wide properties are evaluated independently of individual reference transcripts.

Each query is then compared against all reference transcripts whose genomic span overlaps it. References on the opposite strand are set aside as antisense candidates and are used only if no same-strand evidence is found. Each same-strand reference yields one of six evidence classes, ordered by decreasing specificity. When the query and the reference share an identical junction chain, the pair is full-splice match (FSM); when the query’s junction chain is a contiguous sub-chain of the reference’s, the pair is incomplete-splice-match (ISM). Otherwise the pair is assigned as sharing at least one complete junction, sharing at least one splice-site coordinate, or merely sharing exon sequence, in that order; non-overlapping pairs are discarded. Mono-exon transcripts are handled separately: a mono-exon query overlapping a mono-exon reference is treated as FSM, a mono-exon query contained within a reference exon as ISM, and a mono-exon query spanning a known intron as novel-in-catalog (NIC).

A query transcript may overlap multiple reference transcripts or even multiple genes. IsoMatch ranks these comparisons and identifies the best hit after pairwise comparison. The best hit within each overlapping gene is first selected, followed by selection of the best hit across genes, when applicable. Within a gene, candidates are first compared by evidence class, and only candidates within the same class are ranked further. For FSM and ISM candidates, smaller differences in TSS and TES are preferred. Candidates sharing at least one complete junction are ranked first by the number of matched splice sites, then by exonic overlap in bases, and finally by the smallest difference in exon count. All remaining candidates are ranked using a composite structural score defined as the number of matched splice sites, plus exonic overlap normalized by query length, plus genomic overlap normalized by reference length, minus the absolute difference in exon count.

Candidates that reach only splice-site- or junction-level agreement are temporarily classified as novel-not-in-catalog (NNC) and are further refined using catalog-level evidence. A query associated with a single gene is reclassified as NIC if all of its splice sites are annotated, reflecting a novel combination of known elements. A query containing any unannotated splice site remains NNC. Queries associated with more than one gene are classified as fusion, except when at least one query junction is shared by multiple associated genes. In this case, the ambiguity is attributed to overlapping annotation rather than a genuine fusion event, and the transcript is labeled accordingly. For these transcripts, the reported gene identifier is the concatenation of all associated genes.

#### Output of IsoMatch classify

Classification results are written to two output files. The first is a tabular file in which each row represents a query transcript, with its classification and associated reference transcript information provided in the corresponding columns. The second is a gzip-compressed GTF containing the query transcript models with classification results added as attributes. These attributes include the assigned reference transcript (ISOM_REF_TX_ID), the associated reference gene identifier and gene name (ISOM_REF_GENE_ID and ISOM_REF_GENE_NAME), and the assigned structural category and subcategory (ISOM_CATEGORY and ISOM_SUBCATEGORY).

### 6. Benchmarking

#### WTC11 dataset preprocessing

The WTC11 datasets used for benchmarking were downloaded from the LRGASP project. Raw reads were mapped to the GRCh38 reference genome using platform-specific parameters. Minimap2(Li 2018) (2.28-r1209) was used for read alignment with the LRGASP reference genome, which is based on GRCh38, using platform-dependent parameters (PacBio: -ax splice:hq -uf; ONT: -ax splice -uf-k14). IsoQuant (3.10.0) was then used with the LRGASP reference annotation to detect transcripts in each dataset using the following command:

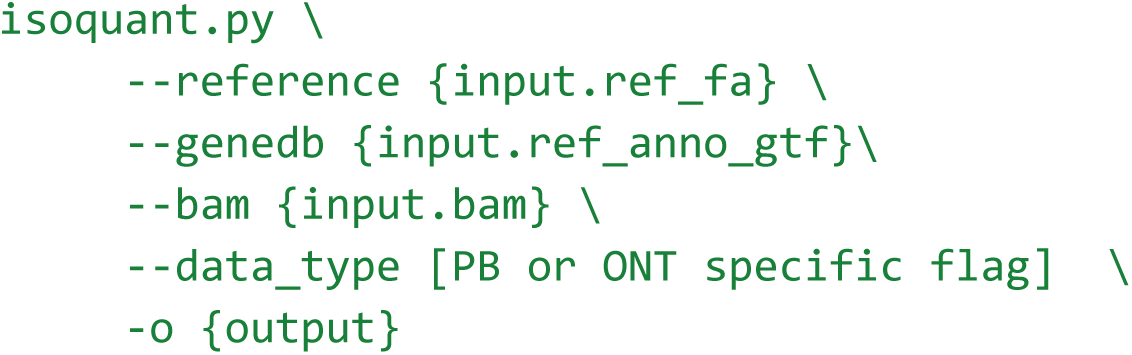

Transcript callsets from each GTF were further divided into multi-exon transcript callsets and mono-exon callsets.

#### Multi-exon transcript evaluation

Unique junction chains (UJCs) were used to compare each tool’s pre-merge input and post-merge output. UJCs were generated with analysis/gtf2ujc.py from the IsoMatch GitHub repository, which ignores mono-exon transcripts and defines each UJC by chromosome, strand, and the ordered intron chain using 1-based inclusive intron coordinates. Merged multi-exon outputs from each tool were converted to UJC tables in the same way and compared with the input UJC set. Retained UJCs were defined as input UJCs present in the merged output. Lost-UJCs were input UJCs absent from the merged output. Novel-UJCs were output UJCs absent from the input. UJCs represented by multiple output transcripts were identified, and the corresponding terminally distinct transcripts were examined and labeled as terminally distinct transcripts sharing the same UJC.

Transcript callsets were merged using IsoMatch (v0.6.0), StringTie (v3.0.3), GffCompare (v0.12.9), and TAMA (commit 2fa3c30) with the following commands.

IsoMatch

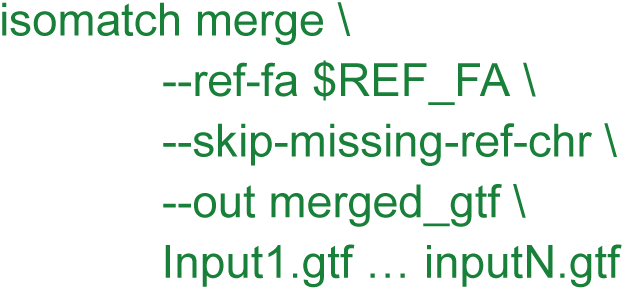

StringTie

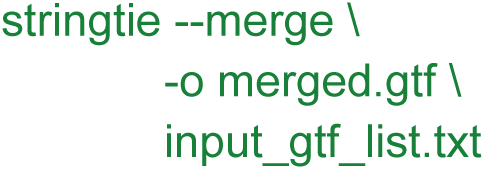

GffCompare

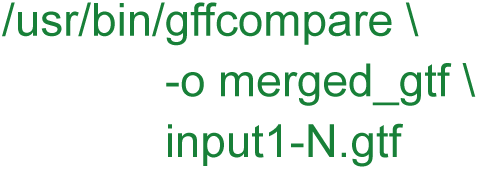

TAMA

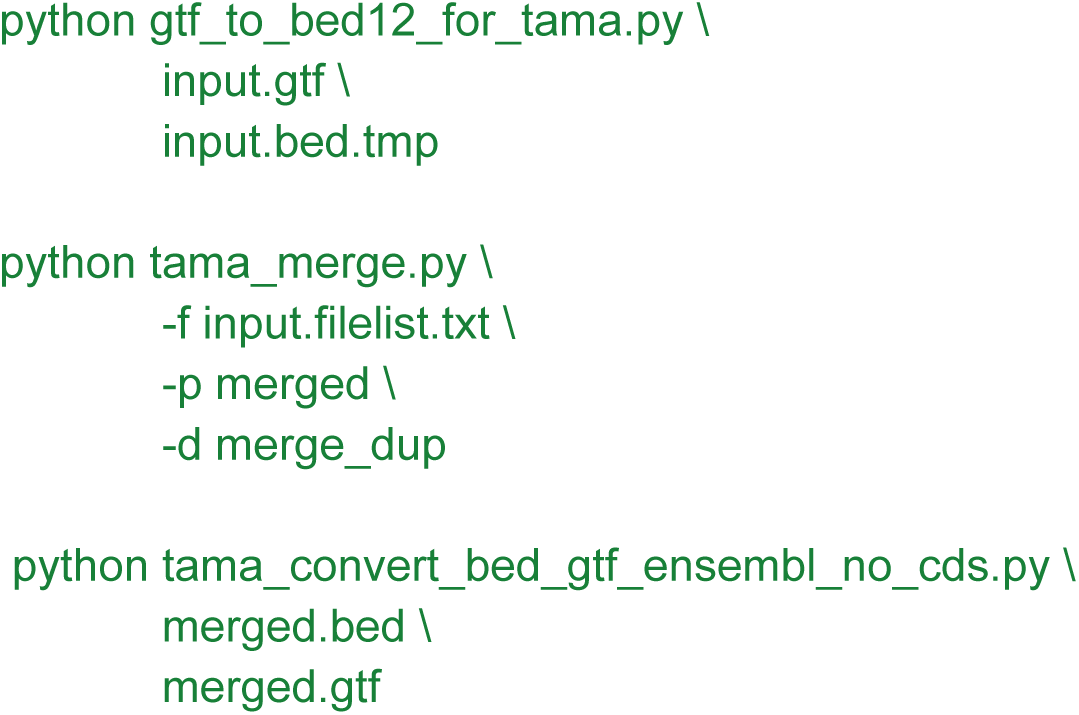

For sequencing protocols using CapTrap, capped was specified in input.filelist.txt; for all other protocols, no_cap was used.

For the strict splice-junction-only comparison, parameters were chosen to minimize terminal-boundary effects. IsoMatch was run with splice-site wobble disabled and terminal refinement disabled for both canonical and non-canonical junctions: -d 0, -a 0, -u 0, -t none, -D 0, -A 0, -U 0, and -T none. StringTie was run in --merge mode with the input list, and the additional permissive filters -c 0, -F 0, -T 0, -i and -f 0. TAMA was run with the same -d merge_dup duplicate-merging mode, but all input BED files were treated as capped with equal priority, and the additional settings -m 0, -a 1000000000, -z 1000000000, and -e common_ends were used so that transcripts sharing the same splice junction chain could be merged without being separated by 5’ or 3’ terminal differences. GffCompare was changed from the default direct-input run to an input-list run with -i, and -D. These settings were used only to test whether each tool could preserve the UJC set under junction-only conditions.

Terminal-support analyses used external human GRCh38 TSS and TES evidence BED files derived from refTSS (IsoMatch GitHub repo: evidence/human.grch38.tss.bed) and PolyASite (IsoMatch GitHub repo: evidence/human.grch38.tes.bed), respectively. To account for 1 bp evidence intervals and small terminal-coordinate differences, each transcript TSS and TES interval was expanded by 10 bp on both sides before matching. Supported transcript IDs were counted with strand-specific intersections using bedtools(Quinlan and Hall 2010) with parameters intersect -u -s. TSS support, TES support, and support at both ends were reported as counts and fractions.

For IsoMatch guide mode, the same multi-exon filtered inputs were merged with --guide-tss and --guide-tes using the refTSS and PolyASite evidence BED files. The guided and default IsoMatch outputs were compared by UJC set changes and by TSS/TES evidence support described above. For terminally distinct transcripts sharing the same UJC retained by IsoMatch and TAMA, transcript IDs sharing the same output UJC were extracted from the UJC tables, terminal sites were computed from the corresponding GTFs, and the same +/-10 bp strand-aware refTSS/PolyASite intersection was used to quantify whether terminally distinct duplicate records were supported by external evidence.

#### Mono-exon transcript evaluation

The filtered mono-exon GTFs were merged by tools using IsoMatch, StringTie merge, TAMA merge, and GffCompare with the -C option. Mono-exon terminal sites were then extracted from the merged GTFs, excluding any output transcript with more than one exon. TSS and TES support was assessed with the same +/- 10 bp strand-aware intersection against refTSS and PolyASite evidence.

#### Synthetic stress test with 1,000 inputs for transcript merging

The GENCODE v49 basic GTF, containing approximately 280,000 transcripts, was supplied 1,000 times as input to the merge command. The commands were:

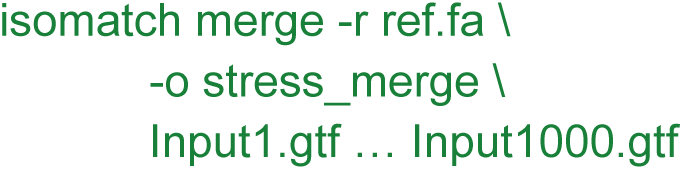
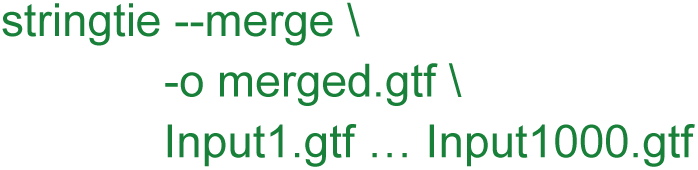

#### Evaluation of transcript classification

We evaluated the classification performance of IsoMatch and compared it with SQANTI3 (6.0.1). The merged transcript set generated in the previous step was used as input for both tools. For IsoMatch, the following command was used:

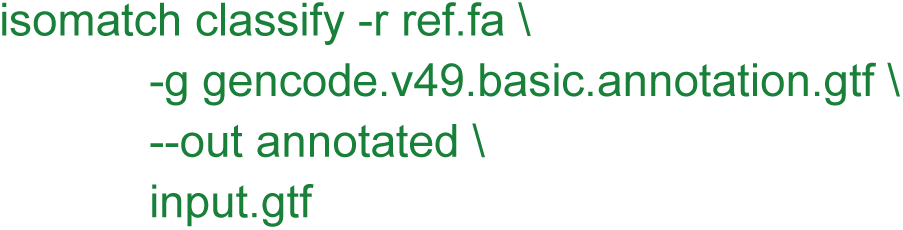

SQANTI3:

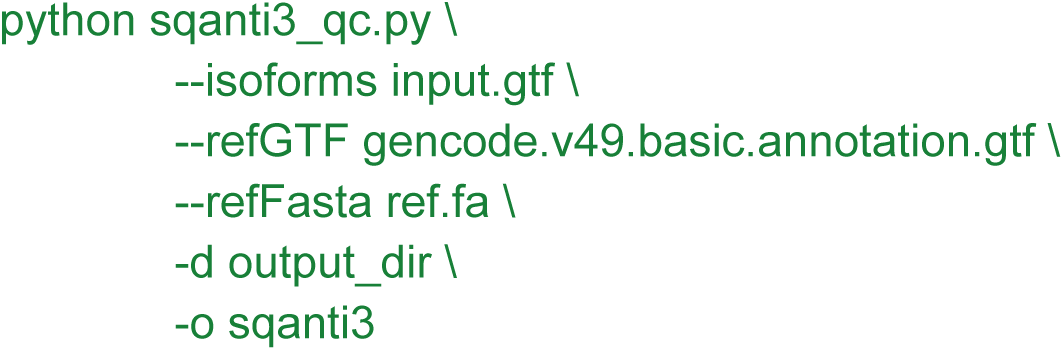

The outputs from IsoMatch and SQANTI3 were parsed, and the structural categories and subcategories were extracted for consistency analysis.

#### Evaluation of computational performance

We downloaded 163 GTF files from the ENCODE project to evaluate the CPU runtime and memory usage of IsoMatch and other transcript-merging tools, including StringTie, GffCompare, and TAMA. All tools were run with their default parameters using the commands described above. Performance was evaluated with 10, 20, 50, 100, and all 163 input GTF files. For the merging benchmark, IsoMatch indexes were built in advance. Each index only needs to be built once and can be reused across runs. Index construction can also be parallelized across input files, with its CPU runtime depending on the degree of parallelization. We therefore excluded index construction from the reported merging runtime. For classification performance, IsoMatch and SQANTI3 were evaluated using the merged GTF generated by IsoMatch as input. The /usr/bin/time utility was used to record CPU runtime and peak memory usage for both the merging and classification steps. Benchmarks were run on a workstation with 64 GB of RAM and an AMD Ryzen 9 7940HX CPU (16 cores, 32 threads).

#### Reference annotation merging and comparison

Reference annotations, including GENCODE v49 Basic, RefSeq v110, and CHESS v3.1.3, were downloaded in GRCh38 coordinates. Only transcripts located on the main chromosomes were retained. Chromosome names were standardized to UCSC-style naming across all annotations. IsoMatch was then used to merge the reference annotations using the following command:

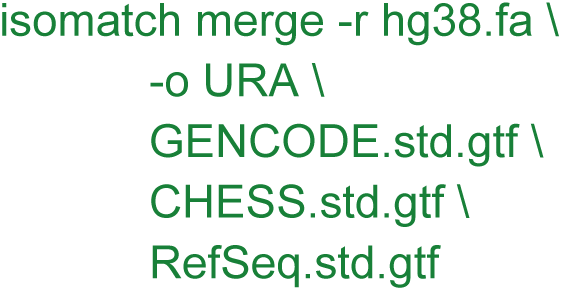

The URA.present_absent.tsv.gz file was parsed to determine the overlap relationships among the input annotations. The merged GTF was used as the URA for downstream analyses.

To evaluate the effects of TSS and TES on reference transcript merging, we tested different configurations that considered splice junctions alone or in combination with TSS and TES. The default configuration considers splice junctions together with both TSS (50 bp wobble) and TES (50 bp wobble). The SJ-only configuration used -t none -T none to disable terminal refinement for both canonical and non-canonical transcripts. The SJ + TSS configuration used -t tss -T tss, whereas the SJ + TES configuration used -t tes -T tes.

#### MANE Select transcript analysis

MANE Select transcripts were identified in the RefSeq GTF by the presence of tag “MANE Select” in their transcript attributes. Corresponding Ensembl transcript IDs were obtained from the db_xref attribute (e.g., db_xref “Ensembl:ENST00000641515.2”), and only transcripts linked to an ENST ID were included in downstream analyses. Merging relationships were traced using the unified reference annotation (URA) generated in the previous step. Specifically, the ISOM_SRC attribute was parsed to recover the original transcript IDs and their source files. Additional non-MANE transcripts assigned to groups containing MANE Select transcripts were identified and characterized according to their structural differences from the corresponding MANE transcripts.

#### Analysis of 119 1000 Genomes project transcript sets

Raw reads were downloaded from the 1000 Genomes Project FTP site and mapped to the GRCh38 reference genome using minimap2 (v2.28-r1209) with PacBio-specific parameters (-ax splice:hq -uf). The resulting alignments were sorted and indexed using SAMtools(Danecek et al. 2021) (v1.16). StringTie (v3.0.3) was then used to assemble transcripts in unguided mode with the following command:

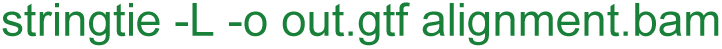

Output GTF files were merged using Isomatch with command:

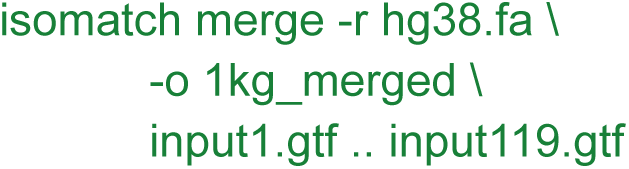

The merged GTF was subsequently classified with GENCODE, RefSeq and CHESS and the URA generated by the previous step. The commands are:

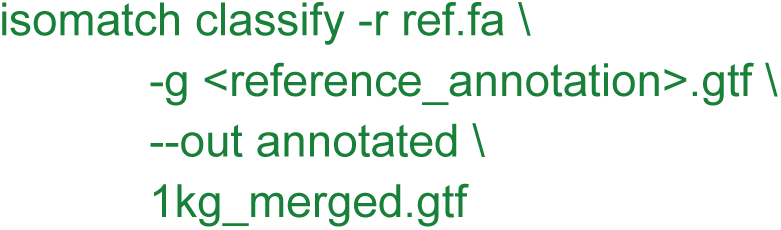

The classification and merge outputs were parsed together to determine the structural category assigned under each reference annotation. Population frequency was extracted from the ISOM_SAMPLE_FREQ attribute in the merged GTF.

## Data availability

WTC11 datasets were downloaded from the LRGASP website: https://www.gencodegenes.org/pages/LRGASP/. GENCODE annotation was downloaded from https://www.gencodegenes.org/human/releases.html. NCBI RefSeq annotation was downloaded from https://ftp.ncbi.nlm.nih.gov/refseq/H_sapiens/annotation/GRCh38_latest/refseq_identifiers/ CHESS annotation was downloaded from https://github.com/chess-genome/chess.

ENCODE GTFs were downloaded from https://www.encodeproject.org/, the accessions are available at **Table S7**. The 1000 Genomes Project long-read RNA datasets were downloaded from https://s3.amazonaws.com/1000g-ont/index.html?prefix=Kinnex/kinnex_1KGP/data/raw/, the accessions are available at **Table S7**. refTSS (v4.1) dataset was downloaded from http://reftss.clst.riken.jp/datafiles/current/human/refTSS_v4.1_human_coordinate.hg38.bed.txt.gz. PolyAsite (v2.0) dataset was downloaded from https://polyasite.unibas.ch/download/atlas/2.0/GRCh38.96/atlas.clusters.2.0.GRCh38.96.bed.gz.

## Code availability

The IsoMatch source code and binaries are available at https://github.com/zhengxinchang/isomatch

## Acknowledgements

The authors would like to thank Zev Kronenberg for helpful discussions. This research was supported in part by the National Institutes of Health (1R01HG011774-01A1, 1U01HG011758-01). We thank the 1000 Genomes Project Long-read Sequencing Consortium (1KGP-LRSC), led by Danny E. Miller and Evan E. Eichler, for generating and publicly releasing the long-read RNA sequencing data used in this study, and PacBio for contributing the Kinnex full-length RNA sequencing. We acknowledge the 1000 Genomes Project and the International Genome Sample Resource (IGSR) for sample and data resources, and the NHGRI Sample Repository for Human Genetic Research at the Coriell Institute for Medical Research for providing the cell lines.

## References

1000 Genomes Project Consortium, Gonçalo R. Abecasis, David Altshuler, et al. 2010. “A Map of Human Genome Variation from Population-Scale Sequencing.” Nature 467 (7319): 1061–1073.

1000 Genomes Project Consortium, Adam Auton, Lisa D. Brooks, et al. 2015. “A Global Reference for Human Genetic Variation.” Nature 526 (7571): 68–74.

Abugessaisa, Imad, Shuhei Noguchi, Akira Hasegawa, et al. 2019. “RefTSS: A Reference Data Set for Human and Mouse Transcription Start Sites.” Journal of Molecular Biology 431 (13): 2407–2422.

Ament, Isabelle Heifetz, Nicole DeBruyne, Feng Wang, and Lan Lin. 2025. “Long-Read RNA Sequencing: A Transformative Technology for Exploring Transcriptome Complexity in Human Diseases.” Molecular Therapy: The Journal of the American Society of Gene Therapy 33 (3): 883–894.

Clavell-Revelles, Pau, Fairlie Reese, Sílvia Carbonell-Sala, et al. 2025. “Long-Read Transcriptomics of a Diverse Human Cohort Reveals Ancestry Bias in Gene Annotation.” Nature Communications 16 (1): 10194.

Danecek, Petr, James K. Bonfield, Jennifer Liddle, et al. 2021. “Twelve Years of SAMtools and BCFtools.” GigaScience 10 (2): giab008.

Glinos, Dafni A., Garrett Garborcauskas, Paul Hoffman, et al. 2022. “Transcriptome Variation in Human Tissues Revealed by Long-Read Sequencing.” Nature 608 (7922): 353–359.

Goldfarb, Tamara, Vamsi K. Kodali, Shashikant Pujar, et al. 2025. “NCBI RefSeq: Reference Sequence Standards through 25 Years of Curation and Annotation.” Nucleic Acids Research 53 (D1): D243–D257.

GTEx Consortium. 2013. “The Genotype-Tissue Expression (GTEx) Project.” Nature Genetics 45 (6): 580–585.

Hamaguchi, Yu, Chao Zeng, and Michiaki Hamada. 2021. “Impact of Human Gene Annotations on RNA-Seq Differential Expression Analysis.” BMC Genomics 22 (1): 730.

Herrmann, Christina J., Ralf Schmidt, Alexander Kanitz, Panu Artimo, Andreas J. Gruber, and Mihaela Zavolan. 2020. “PolyASite 2.0: A Consolidated Atlas of Polyadenylation Sites from 3’ End Sequencing.” Nucleic Acids Research 48 (D1): D174–D179.

Jetzinger, Fabian, Alejandro Paniagua, Stanley Cormack, et al. 2026. “Handling Biological Replicates in Long-Read RNA Sequencing Data by Joining or Not Joining.” Nature Communications, ahead of print, September 1. 10.1038/s41467-026-77367-6.

Kouam, Cedric, Jackson Mingle, Pilar Álvarez Jerez, et al. 2026. “SALRR: Scalable Analysis of Long-Read RNA-Seq Enables Comprehensive Transcriptome Profiling in Human Brain.” In bioRxiv. BioRxiv, August 29. 10.64898/2026.08.27.747499.

Kovaka, Sam, Aleksey V. Zimin, Geo M. Pertea, Roham Razaghi, Steven L. Salzberg, and Mihaela Pertea. 2019. “Transcriptome Assembly from Long-Read RNA-Seq Alignments with StringTie2.” Genome Biology 20 (1): 278.

Kuo, Richard I., Yuanyuan Cheng, Runxuan Zhang, et al. 2020. “Illuminating the Dark Side of the Human Transcriptome with Long Read Transcript Sequencing.” BMC Genomics 21 (1): 751.

Li, Heng. 2018. “Minimap2: Pairwise Alignment for Nucleotide Sequences.” Bioinformatics (Oxford, England) 34 (18): 3094–3100.

Lucas, Julian K., Prajna Hebbar, Wen-Wei Liao, et al. 2026. “HPRC2: A Human Pangenome Reference with near-Complete Coverage of Common Genetic Variation.” In bioRxivorg. BioRxiv, July 22. 10.64898/2026.07.21.739710.

Monzó, Carolina, Tianyuan Liu, and Ana Conesa. 2025. “Transcriptomics in the Era of Long-Read Sequencing.” Nature Reviews. Genetics 26 (10): 681–701.

Morales, Joannella, Shashikant Pujar, Jane E. Loveland, et al. 2022. “A Joint NCBI and EMBL-EBI Transcript Set for Clinical Genomics and Research.” Nature 604 (7905): 310–315.

Mudge, Jonathan M., Sílvia Carbonell-Sala, Mark Diekhans, et al. 2025. “GENCODE 2025: Reference Gene Annotation for Human and Mouse.” Nucleic Acids Research 53 (D1): D966–D975.

Pardo-Palacios, Francisco J., Angeles Arzalluz-Luque, Liudmyla Kondratova, et al. 2024. “SQANTI3: Curation of Long-Read Transcriptomes for Accurate Identification of Known and Novel Isoforms.” Nature Methods 21 (5): 793–797.

Pardo-Palacios, Francisco J., Dingjie Wang, Fairlie Reese, et al. 2024. “Systematic Assessment of Long-Read RNA-Seq Methods for Transcript Identification and Quantification.” Nature Methods 21 (7): 1349–1363.

Patowary, Ashok, Pan Zhang, Connor Jops, et al. 2024. “Developmental Isoform Diversity in the Human Neocortex Informs Neuropsychiatric Risk Mechanisms.” Science (New York, N.Y.) 384 (6698): eadh7688.

Pertea, Geo, and Mihaela Pertea. 2020. “GFF Utilities: GffRead and GffCompare.” F1000Research 9 (April): 304.

Pertea, Mihaela, Geo M. Pertea, Corina M. Antonescu, Tsung-Cheng Chang, Joshua T. Mendell, and Steven L. Salzberg. 2015. “StringTie Enables Improved Reconstruction of a Transcriptome from RNA-Seq Reads.” Nature Biotechnology 33 (3): 290–295.

Pertea, Mihaela, Alaina Shumate, Geo Pertea, et al. 2018. “CHESS: A New Human Gene Catalog Curated from Thousands of Large-Scale RNA Sequencing Experiments Reveals Extensive Transcriptional Noise.” Genome Biology 19 (1): 208.

Prjibelski, Andrey D., Alla Mikheenko, Anoushka Joglekar, et al. 2023. “Accurate Isoform Discovery with IsoQuant Using Long Reads.” Nature Biotechnology 41 (7): 915–918.

Quinlan, Aaron R., and Ira M. Hall. 2010. “BEDTools: A Flexible Suite of Utilities for Comparing Genomic Features.” *Bioinformatics (Oxford*, England) 26 (6): 841–842.

Reese, Fairlie, Brian Williams, Gabriela Balderrama-Gutierrez, et al. 2023. “The ENCODE4 Long-Read RNA-Seq Collection Reveals Distinct Classes of Transcript Structure Diversity.” In bioRxivorg. May 16. 10.1101/2023.05.15.540865.

Reyes, Alejandro, and Wolfgang Huber. 2018. “Alternative Start and Termination Sites of Transcription Drive Most Transcript Isoform Differences across Human Tissues.” Nucleic Acids Research 46 (2): 582–592.

Shinder, Ida, Geo Pertea, Richard Hu, Zoe Rudnick, and Mihaela Pertea. 2026. “StringTie3 Improves Total RNA-Seq Assembly by Resolving Nascent and Mature Transcripts.” Nature Methods 23 (6): 1126–1137.

Tushev, Georgi, Caspar Glock, Maximilian Heumüller, Anne Biever, Marko Jovanovic, and Erin M. Schuman. 2018. “Alternative 3’ UTRs Modify the Localization, Regulatory Potential, Stability, and Plasticity of mRNAs in Neuronal Compartments.” Neuron 98 (3): 495–511.e6.

Varabyou, Ales, Markus J. Sommer, Beril Erdogdu, et al. 2023. “CHESS 3: An Improved, Comprehensive Catalog of Human Genes and Transcripts Based on Large-Scale Expression Data, Phylogenetic Analysis, and Protein Structure.” Genome Biology 24 (1): 249.

Wang, Eric T., Rickard Sandberg, Shujun Luo, et al. 2008. “Alternative Isoform Regulation in Human Tissue Transcriptomes.” Nature 456 (7221): 470–476.

Wang, Xi, Jingyi Hou, Claudia Quedenau, and Wei Chen. 2016. “Pervasive Isoform-Specific Translational Regulation via Alternative Transcription Start Sites in Mammals.” Molecular Systems Biology 12 (7): 875.

Zheng, Xinchang, Sonia Garcia-Ruiz, Emil K. Gustavsson, Mina Ryten, and Fritz J. Sedlazeck. 2026. “The Novel Transcripts We Keep Rediscovering.” Nature Biotechnology, ahead of print, September 22. 10.1038/s41587-026-03321-y.

