## supplementary Notes for "IsoMatch enables population-scale transcript integration and reveals reference-dependent isoform novelty"

### Supplementary Note

#### Note 1: evaluation of the impact of junction wobble tolerance

IsoMatch applies a default splice-junction wobble tolerance of 3 bp to transcripts containing at least one non-canonical splice junction. This design aims to reduce potentially artifactual transcript models while preserving well-supported splice structures. As described in the Methods, IsoMatch first merges transcripts whose splice junctions are all canonical (GT-AG, GC-AG, or AT-AC). It then assigns non-canonical transcripts to compatible canonical groups when their splice junctions match within the specified tolerance. The rationale is that a non-canonical junction close to a canonical junction may reflect a small positional error rather than a distinct biological splice event. Non-canonical transcripts without a compatible canonical group are subsequently merged among themselves using the same tolerance, allowing potentially genuine non-canonical structures to remain represented in the output.
To quantify the effect of this tolerance, we jointly merged all 18 WTC11 samples while varying the non-canonical donor (`-D`), acceptor (`-A`), and unstranded splice-junction (`-U`) wobble tolerances together from 0 to 10 bp, with canonical splice-junction tolerances fixed at 0 bp. The number of merged transcript models decreased modestly from 127,977 at 0 bp to 122,225 at 10 bp (**Supplementary Note Figure S1**). At the default tolerance of 3 bp, IsoMatch produced 125,010 transcript models, 2,967 fewer than at 0 bp. At this setting, 3,287 non-canonical source transcripts belonged to 2,207 merged groups that also contained canonical transcripts. We extracted these source transcripts with their original structures and classified them against the integrated reference annotation (URA). Most were classified as NNC (93.55%), whereas 3.10% were FSM (**Supplementary Note Table S1**). These results show that the non-canonical source transcripts incorporated into canonical groups predominantly contained splice junctions absent from the reference annotation. Although this pattern is consistent with the intended consolidation of potentially artifactual models, reference-based classification alone cannot distinguish technical errors from genuine non-canonical splicing or establish whether individual merges were erroneous. The wobble tolerances are fully user-adjustable, allowing users to choose the degree of consolidation appropriate for their analysis.

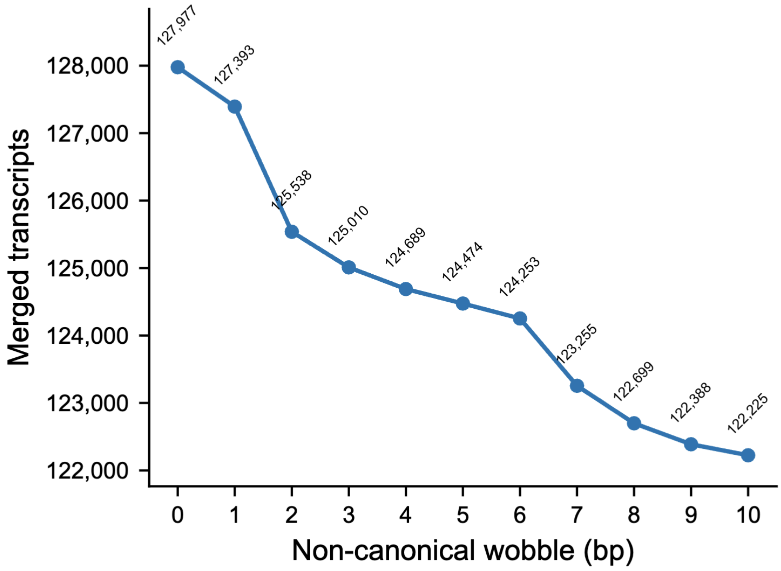


***Supplementary Note Figure S1.*** *Effect of non-canonical splice-junction wobble on transcript merging. All 18 WTC11 samples were merged jointly using IsoMatch with non-canonical wobble tolerances ranging from 0 to 10 bp and canonical wobble tolerances fixed at 0 bp. The number of merged transcript models is shown for each setting; the default non-canonical tolerance is 3 bp.*

**Supplementary Note Table S1.** Classification of non-canonical source transcripts incorporated into canonical-containing merge groups. At a non-canonical wobble tolerance of 3 bp, 3,287 non-canonical source transcripts were identified in 2,207 merged groups containing canonical transcripts. Their original, pre-merge structures were classified against the URA. Percentages are calculated relative to the 3,287 source transcripts.

| **classification** | **count** | **Percent of all 3,287 transcripts** |
|---|---|---|
| FSM | 102 | 3.10% |
| ISM | 0 | 0% |
| NIC | 33 | 1.00% |
| NNC | 3,075 | 93.55% |
| Other | 77 | 2.34% |

#### Note 2: Differences between classification strategy of IsoMatch and SQANTI3

There were 2,992 discordant transcripts (2.39%) between IsoMatch and SQANTI3. This note examines six category-pair combinations totaling 2,776 transcripts (92.78% of discordant calls; 2.22% of the catalog). The full discordance table is available at Supplementary **Table S4**:

***Supplementary Note Table S2****: Major discordant structural-category assignments between IsoMatch and SQANTI3.*

| Transcript count | IsoMatch category | SQANTI3 category |
|---|---|---|
| 1,356 | ISM | NIC |
| 679 | Genic | NNC |
| 123 | Genic | antisense |
| 63 | Genic | fusion |
| 26 | Genic | NIC |
| 529 | antisense | Genic intron |

**IsoMatch ISM versus SQANTI3 NIC**
This discordance involved 1,356 transcripts on both strands (634 on the plus strand and 722 on the minus strand). In every case, the genomic leftmost terminal exon spanned at least one intron of the reference transcript selected by IsoMatch, while the query splice-junction chain formed a contiguous subset of the reference junction chain. Among these transcripts, 60 also had a genomic rightmost terminal exon spanning a reference intron. Only 1,057 spanned the reference transcript's genomic leftmost intron; therefore, these events cannot collectively be described as first-intron retention.

IsoMatch determines ISM status based on the contiguous junction-chain subset relationship and subsequently evaluates intron-retention subtypes, classifying all 1,356 transcripts as ISM with intron retention. In contrast, SQANTI3/Cupcake compares exon boundaries starting from the first overlapping exon in genomic coordinate order. When applied to the same query-reference pairs, its `compare_junctions` function returned partial (primary class for NIC and NNC) for all 1,356 transcripts and never returned subset (primary class for ISM). SQANTI3 subsequently assigned these transcripts to NIC through its known-gene novel classification branch: 1,335 were annotated as intron_retention, whereas the remaining 21 were associated with different genes and annotated as combination_of_known_junctions. Thus, this discrepancy reflects a slightly different implementation of ISM in the presence of terminal intron retention in IsoMatch and SQANTI3.

**IsoMatch genic versus SQANTI3 NNC/antisense/fusion/NIC**
Both tools use genic as a fallback category for transcripts with exonic overlap but insufficient evidence for a splice-based category. However, IsoMatch requires an exact shared junction or qualifying exon boundary before assigning a novel splice category, prioritizes same-strand overlap over antisense evidence, and performs fusion refinement only for transcripts that first enter the NNC pathway. SQANTI3 additionally accepts partial exon-chain matches, retains antisense evidence during mono-exon classification, and can combine multiple gene associations into a fusion assignment.

For the 679 genic-to-NNC transcripts, SQANTI3 accepted partial-like exon-chain matches that did not satisfy the IsoMatch requirements for NNC. The 123 genic-to-antisense transcripts had both same-strand and opposite-strand exonic overlap; IsoMatch prioritized the same-strand overlap, whereas SQANTI3 retained the antisense association. For the 63 genic-to-fusion transcripts, both tools generally identified the same associated genes, but only SQANTI3 promoted the multi-gene evidence to fusion. Finally, 25 of the 26 genic-to-NIC transcripts resulted from conditional opposite-strand handling in the SQANTI3 mono-exon branch, while the remaining transcript reflected a boundary-coordinate difference between the two implementations.

**IsoMatch antisense versus SQANTI3 genic intron**

IsoMatch and SQANTI3 differ in their treatment of antisense transcripts because they collect antisense evidence differently. IsoMatch classifies a transcript as antisense whenever its genomic span overlaps an annotated gene on the opposite strand, prioritizing this evidence over genic-intron and intergenic assignments. In contrast, SQANTI3 may require exon-level overlap in some classification branches; transcripts located entirely within an intron of an opposite-strand gene can therefore lose their antisense association and be classified as genic-intron or intergenic instead. Thus, IsoMatch applies a broader, strand-aware definition of antisense, whereas SQANTI3 uses a more restrictive and context-dependent criterion.
